# Hairpin-RNA Spray Confers Resistance to Mungbean Yellow Mosaic India Virus in Mungbean

**DOI:** 10.1101/2024.03.15.585278

**Authors:** Kiran Vilas Dhobale, Lingaraj Sahoo

## Abstract

The prevalence of Begomovirus diseases poses a significant threat to legume crops, necessitating the exploration of innovative control measures. This investigation explores the utilization of dsRNA molecules to initiate RNA interference (RNAi) targeting begomovirus, particularly focusing on *Mungbean yellow mosaic India virus* (MYMIV) and its potential threat to mungbean crops. Given the lack of genetic resistance in commercially available mungbean varieties, the study endeavors to employ RNAi as a strategic method for the effective control of MYMIV. The approach involves the preparation of vectors for the transient expression of three dsRNA targeting multiple overlapping ORFs of MYMIV DNA A through agroinoculation, and the selection of a highly efficient construct for dsRNA expression in bacteria, enabling topical application to mungbean plants in growth chamber experiments. Agroinoculation assays demonstrate effective resistance against MYMIV, as confirmed by reduced symptom severity, limited virus accumulation, and the presence of viral mRNAs. The stability of the prepared dsRNA against nucleases is confirmed, showcasing its ability to enter plant cells, move to non treated trifoliate leaves, and form siRNA when sprayed onto mungbean leaves, as validated by qRT-PCR and northern blotting. Varied combinations of the timing of dsRNA spray and virus infection reveal differential resistance against the virus. Notably, spraying two days before or on the same day as virus exposure emerges as the most suitable time to achieve optimal resistance against virus infection. In light of these findings, the topical application of dsRNAs stands out as a promising and effective strategy for MYMIV control in mungbean crops.

## INTRODUCTION

Mungbean (*Vigna radiata* L. Wilczek) is an important legume crop in India and South Asian countries, providing an affordable source of high-quality dietary proteins (1,2). Cultivated in tropical and sub-tropical regions globally, India is a prominent producer and consumer of mungbean (3). Despite its significance, the average productivity of mungbean in India remains low (3). Yellow mosaic disease (YMD) poses a major challenge to mungbean cultivation worldwide (4,5). Caused by yellow mosaic viruses (YMVs) transmitted by the whitefly (6,7), YMD leads to crop yield losses ranging from 10 to 100% (8). The disease is prevalent in countries like India, Bangladesh, and Pakistan (4). YMD is caused by four distinct begomoviruses, collectively known as yellow mosaic viruses (YMVs) (9), including *Mungbean Yellow Mosaic Virus* (MYMV) (10), *Mungbean Yellow Mosaic India Virus* (MYMIV) (11), D*olichos Yellow Mosaic Virus* (DoYMV) (12), and *Horsegram Yellow Mosaic Virus* (HgYMV), all belonging to the genus *Begomovirus* within the family *Geminiviridae* (13).

The begomovirus genus, comprising 445 distinct species, is the largest in the virosphere (13). It induces economically significant diseases in crucial crops like mungbean (5). New World (NW) begomoviruses have bipartite genomes, while Old World (OW) begomoviruses exhibit both monopartite and bipartite configurations. The genome structure includes bipartite (DNA-A and DNA-B) or monopartite configurations, each circular single stranded DNA (ssDNA) components being around 2.7 kb (14). Additionally, begomoviruses are associated with circular DNA satellites: betasatellites, alphasatellites, and deltasatellites (15–17). Begomovirus proteins, play multifunctional roles crucial for disease development. DNA-A features six open reading frames (ORFs): two in the virion sense (AV1 and AV2) and four in the complementary sense (AC1, AC2, AC3, and AC4). DNA-B comprises two ORFs: BV1 and BC1. AV2 is unique to Old World bipartite begomoviruses, absent in New World viruses. AV1 and AV2 encode capsid protein (CP) and pre-coat protein, respectively (18). AC1, AC2, and AC3 serve as replication initiator protein (Rep) (19,20), transcription activator protein (TrAP) (21,22), and replication enhancer protein (REn) (23–25), respectively. AC4-encoded protein is essential for symptom production (26,27). DNA-B carries BC1 and BV1 ORFs, functioning as movement protein (MP) and nuclear shuttle protein (NSP) (28), respectively (29,30).

Among various YMD management strategies deployed, RNA interference (RNAi) is a highly effective strategy for developing durable resistance against viral diseases in plants (31). Plants employ post-transcriptional gene silencing (PTGS) to silence or knock down the expression of specific viral genes, conferring resistance (32). This mechanism involves the sequence-specific degradation of viral RNA, achieved by processing double-stranded RNA (dsRNA)/hairpin RNA (hpRNA) or partial overlapping transcripts of DNA viruses into small interfering RNA (siRNA) of approximately 21–24 nucleotides. Dicer-like enzymes facilitate this process (33). The processed siRNA binds to argonaute (AGO) protein and incorporates into the RNA-induced silencing complex (RISC), leading to the degradation of target RNA or viral transcripts with sequence similarity to the siRNA (34–36). Additionally, complementary guide RNA can serve as a primer for RNA-dependent RNA polymerase (RDR), generating secondary siRNA and ensuring the amplification of the siRNA signal (37).

The principle of RNAi has been extensively used to engineer transgenic resistance in plants against viruses (38–43). This involves genetically modifying plants with a segment of nucleotide sequence from the viral genome. Transgenic technology has successfully produced begomovirus-resistant cultivars in various legume crops, including mungbean (41). While stable transgene integration allows efficient RNAi induction against geminiviruses, the broader adoption of Host-Induced Gene Silencing (HIGS) faces challenges such as limited transformation protocols, time-consuming processes, high costs, public concerns about genetically modified organisms (GMOs), stringent regulatory laws, and engineered RNA silencing trait instability (44). To address these challenges, an alternative approach has emerged, involving the induction of RNAi in plants against viruses through the external or topical application of dsRNA/hpRNA derived from the viral genome (33). This non-GMO approach is promising, efficient, and socially acceptable for viral disease management (45).

The use of externally applied naked dsRNA to prevent infection by various plant viruses was first reported by Tennellado and Diaz Ruiz in 2001 (46). Subsequent studies confirmed the efficacy of dsRNA vaccination against several plant RNA viruses (47–53). Although limited, there are reports of dsRNA vaccination against DNA viruses like begomoviruses. For instance, Namgiala et al. utilized dsRNA vaccination to confer protection against the bipartite geminivirus *Tomato leaf curl virus* (ToLCV) and the tripartite RNA virus *Cucumber mosaic virus* (CMV) (54). Recently, the approach was successfully employed in blackgram plants against MYMV (55). Topical dsRNA application also shows promise against monopartite geminiviruses such as Tomato yellow leaf curl virus (TYLCV) (56) and Chilli leaf curl virus (ChiLCV) (57).

In the present study, a non-transgenic approach was employed to induce resistance to YMD in mungbean. Initially, three hpRNAi constructs were transiently expressed in the MYMIV-susceptible mungbean cultivar K851, akin to previous studies (41,58), to evaluate their efficacy in providing protection against MYMIV infection. The most efficient hpRNAi clone was then selected, and from this single construct, highly efficient *in vivo*-produced hpRNA molecules were derived. The study revealed that the spray application of hpRNA derived from DNA A genes significantly provides protection against MYMIV in mungbean. Importantly, this research represents the first demonstration of the effectiveness of exogenously applied hpRNA against YMD in mungbean crops.

## MATERIALS AND METHODS

### Biological Materials and Target Region (TR) selection

Mungbean cultivar (cv.) K851, highly susceptible to MYMIV, was cultivated in plastic pots filled with vermiculite and soil under greenhouse conditions (∼25°C, 16 hours light/8 hours darkness) was used throughout the study. Infectious clones of MYMIV mungbean isolates of Orissa (GenBank Accessions: DNA-A, OK431083 and DNA-B, OK431084) were utilized to induce YMD in the mungbean plants (59).

To achieve broad-spectrum resistance against YMD, a total of 25 representative MYMV and MYMIV DNA-A components, originating from 10 different host plants in 6 countries were selected for multiple sequence alignment using the ClustalW algorithm. The sequences were obtained from the NCBI Viral RefSeq database, ensuring a comprehensive representation of the YMD causing begomovirus diversity. Two RNAi target regions (TRs) were carefully selected based on multiple criteria, including the functional significance of viral proteins, overlapping regions of ORFs, sequence length, and sequence conservation. TR-1, spanning 239 base pairs (bp), and TR-2, also 239 bp in length, encompass highly conserved regions within the viral genome. TR-1 includes the overlapping section of the AC4 and AC1 genes, while TR-2 encompasses the overlapping region of the AC2, AC3, and AC1 genes.

### Development of hpRNAi Constructs

Three hpRNAi constructs (hpTR-1_pART27, hpTR-2_pART27, and hpTR-1+2_pART27) were prepared using high fedility PCR based method. Plasmid DNA containing a full-length DNA-A genomic component of MYMIV mungbean isolate of Orissa (Acc. No. OK431083) was used as a template for the amplification of the viral RNAi target regions using primer set with cloning restriction site (**Table 1**). To prepare hpRNAi cassattes, for TR-1 and TR-2 the PCR amplified target fragments were cloned in sense orientation (XhoI and KpnI) and in antisense orientation (XbaI and ClaI) on either sides of PDK-intron of the intermediate vector, pKANNIBAL (CSIRO, Plant Industry, Canberra, Australia). The constructed clones were named as hpTR-1_pKannibal and hpTR-2_pKannibal respectively. For construction of hpTR-1+2 stack RNAi cassatte, the sense fragments of TR-1 (XhoI and EcoRI) and TR-2 (EcoRI and KpnI) interrupted by 8 nt gap, and antisense fragments of TR-1 (XbaI and BamHI) and TR-2 (BamHI and ClaI) interrupted by 8 nt gaps were cloned on either side of the PDK intron of pKANNIBAL and prepared construct labelled as hpTR-1+2_pKannibal. To prepare the agroinfectious hpRNAi constructs, each hpRNAi caassatte (from hpRNAi_pKannibal clones) under the control of CaMV35S promoter and OCS terminator (as NotI fragments) were subcloned into the plant binary vector, pART27 (supplemental Fig. 4). The hpRNAi clones of pART27 were finally transformed into *Agrobacterium tumefaciens* strain EHA105 by electroporation (25 µF, 200 Ω, 2500 V) in a Gene Pulser XCell (Bio-Rad, USA).

**Table 1.**
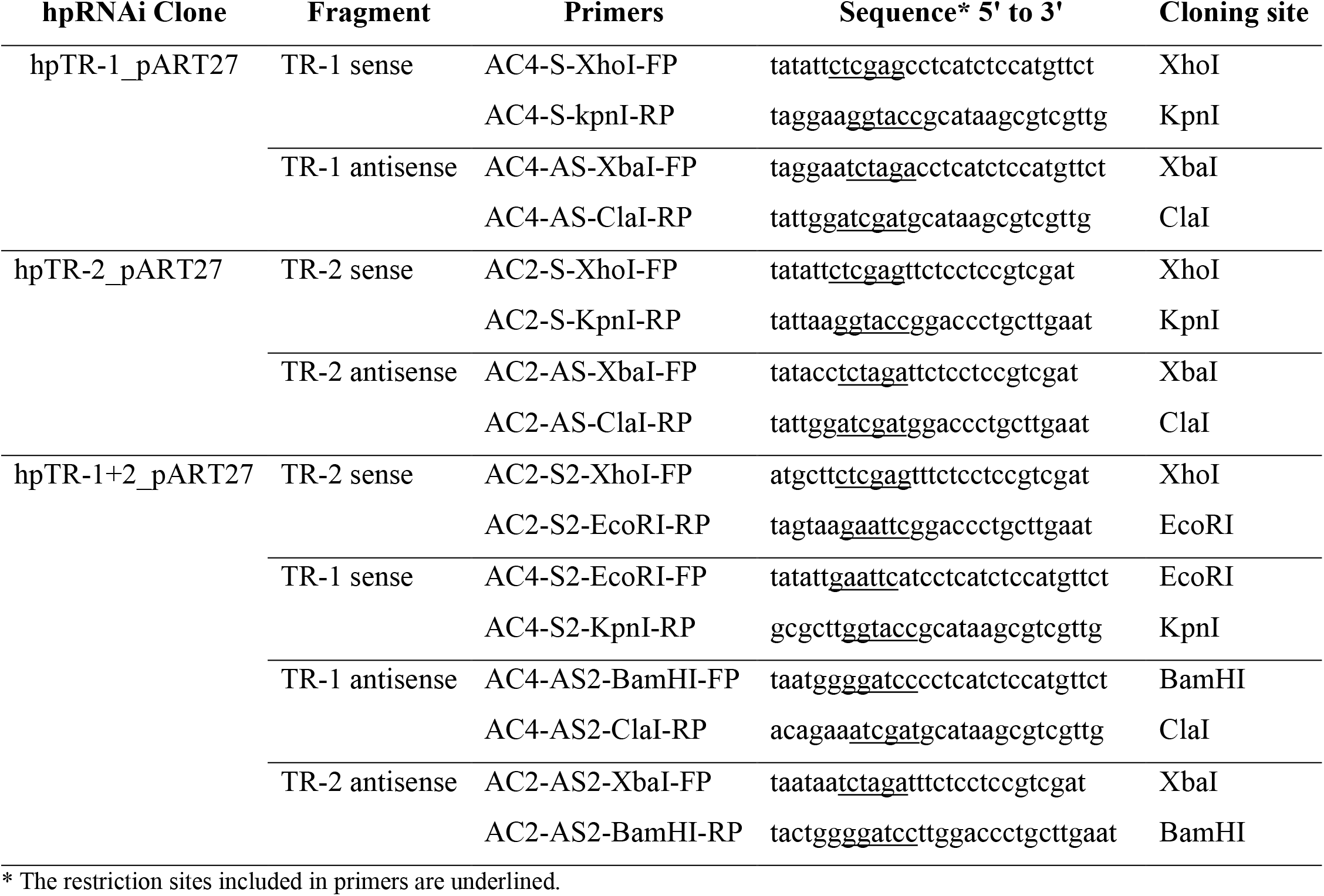
Primers used for amplification of virus sequences for the production of hairpin RNAi constructs.

### Efficacy Validation of hpRNAi Constructs Using Transient Mungbean-MYMIV Pathosystems

*A. tumefaciens* strain EHA105 harbouring hpRNAi_pART27 constructs was used against MYMIV infection in mungbean cv. K851. For transient bioassay about 5-6 plants were infected and three biological replicates were performed. Briefly, the *A. tumefaciens* strain EHA105 harbouring respective clones were cultured separately in YEP medium containing antibiotics (50 μg/ml kanamycin and 20 μg/ml rifampicin) overnight at 28 °C to reach the OD600 = 0.6. Cells were harvested and resuspended in a buffer containing 10 mM 2-(*N-* morpholino) ethanesulfonic acid (MES) and 10 mM MgCl_2_, pH 5.8, and 200 µM acetosyringone. The resuspended cells were agitated at 90 rpm at 28 °C for 1 h before infiltrating the abaxial surface of two trifoliate leaves of 3-4 weeks-old mungbean cv. K851 plants using a 5 ml needleless syringe and the plants were kept in greenhouse at 25 ± 2°C and a 16/8 h light/dark.

For the efficacy validation of three hpRNAi_pART27 clones, the co-infiltration of viral infectious clones (MYMIV 2A + 2B DNA componants) and hpRNAi clones was performed, i.e., an equal volume of each *A. tumefaciens* cultures were mixed prior to agroinoculation. The plants were agroinoculated in five combinations, i.e., 1) MYMIV + hpTR-1_pART27, 2) MYMIV + hpTR-2_pART27, 3) MYMIV + hpTR-1+2_pART27, 4) MYMIV + empty pART27, and 5) empty pART27.

### *In vivo* Production of hpRNA in *E. coli* HT115

To produce hairpin RNA (hpRNA) molecules, the hpRNAi cassette was obtained as an XhoI and XbaI fragment from the hpTR-1+2_pART27 clone. Subsequently, the L4440 vector, harbouring T7 promoters at both ends, was chosen as the recipient vector for subcloning. The hpTR-1+2 cassette was ligated into the L4440 vector, creating the recombinant vector labelled as hpTR-1+2_L4440 clone. This recombinant vector was then transformed into *E. coli* HT115 (DE3) cells, which possess an IPTG-inducible T7 RNA polymerase gene and lack RNAse III activity due to disrupted gene by a Tn*10* transposon (60).

Following the methodology described by (61) with minor modifications (62), single colonies of *E. coli* HT115 transformants carrying the hpTR-1+2_L4440 plasmid were cultured at 37°C with shaking for 16 hours in LB medium supplemented with tetracycline (12.5 µg/mL) and ampicillin (100 µg/mL). Each culture was then diluted 1:100 in a final volume of 0.5 L of LB medium supplemented with the same antibiotics and incubated at 37°C until reaching an OD600 of 0.5. Subsequently, 0.4 mM IPTG was added to induce T7 polymerase expression for an additional 3-4 hours. The cells were harvested by centrifugation, lysed using a lysis buffer (0.1% SDS in 1x PBS), and treated with RNaseA solution (1 μg of RNase enzyme in 5 mM EDTA, 300 mM sodium acetate, 10 mM Tris–Cl pH 8) at 37°C for 30 minutes to degrade single-stranded RNAs. Finally, dsRNA was extracted using TRIzol™ reagent (Invitrogen, USA, Cat. No. 15596026) following the manufacturer’s protocol. The double stranded nature of RNA was validated by incubating hpRNA molecules with DNase I and RNase A.

### Life Span and Systemic Movement of hpRNA and siRNA

In order to evaluate the life-span of hpRNAs and siRNA in the plants, that were not infected with MYMIV, 30 µg of hpRNAs (hpTR-1+2) was mixed with 1 ml of sterile water and mechanically inoculated in two fully expanded trifoliate leaves (1 ml per plant). Five plants were used for each time point. Just before sampling, leaves were washed with Triton X-100 (0.05%) and water to eliminate residual hpRNAs present on the leaf surface. Treated plants were kept in the growth chamber as mentioned above. Trifoliate leaves from treated (local) (at time points 3, 6, 9, 12 dpi) and from non-treated (systemic) leaves (at time points 3, 6, 9, 24 dpi) were collected from all plants.

To assess the internalization of hpRNA within plant cells and its subsequent systemic movement, semi-quantitative reverse transcriptase PCR (semi-qRT-PCR) was performed. Total RNA was isolated from collected leaves using TRIzol reagent, and its quality and concentration were assessed. cDNAs were generated from 50 ng of total RNA using gene-specific primer pair AC2_S2_xhoI_FP and AC4_S2_kpnI_RP **(Table 1)**. This same primer pair was employed to identify the presence of hpRNA (for hpTR_1+2) via PCR in both local and systemic leaves.

For the detection of hpRNA conversion to siRNA and its systemic movement, a northern blot analysis was carried out. Total RNA was isolated using TRIzol extraction and enriched for low molecular weight (LMW) RNAs in line with the procedure described by Peng et al. (63). Subsequently, 15 μg of LMW RNA samples underwent electrophoresis in an 18% polyacrylamide (19:1) gel containing 7 M urea and buffered with 0.5 X TBE using an SE600 standard dual-cooled gel electrophoresis unit (GE Healthcare, USA) until the bromophenol blue dye reached the bottom of the gel. Blots were transferred using a trans-blot SD semi-dry electrophoretic transfer unit (Bio-Rad, Cat. No. 170-3940) onto a Hybond–N membrane (Roche). The membrane was auto-crosslinked at 120,000 μJ in a Stratalinker 1800 (Agilent Technologies, Belgium). Following prehybridization in DIG Easy Hyb hybridization solution (Roche Diagnostics, Belgium) at 65°C for 30 minutes, the membrane was hybridized with a DIG-labeled TR-1+2 RNA probe (spanning 479 nucleotides) at 65°C for 12 hours in a hybridization oven. The DIG-labeled RNA probe was prepared through PCR amplification with M13-FP and AC4-S2-KpnI-RP primers **(Table 1)** using the hpTR-1+2_L4440 plasmid DNA as the template. After purification, 1 μg of the PCR product was used for RNA probe labeling with T7 RNA polymerase as per the DIG Northern Starter Kit’s Instruction Manual (Roche). Post-hybridization washes and immuno-chemiluminescent detection of the bound probe were conducted following the instructions provided in the DIG Northern Starter Kit manual (Roche). To assess siRNA size via northern blot analysis, three oligos (24 nt, 22 nt, and 20 nt) with sequences complementary to the hpRNA probe were commercially synthesized and used as an RNA ladder **(Fig. 4 A**).

### hpRNA Spray Assay

The study aimed to evaluate the effectiveness of hpTR-1+2 against MYMIV through five treatment combinations. These treatments (T1-T5) comprised: (T1) positive control involving MYMIV agroinoculation into three-week-old mungbean cv. K851 plants without hpRNA; (T2) mock inoculation with *A. tumefaciens* carrying empty pCAMBIA3300, followed by a spray of 30 µg of hpTR-1+2; (T3) MYMIV infection, immediately followed by hpTR-1+2 spray on the same day; (T4) topical spray after MYMIV agroinoculation, with hpTR-1+2 sprayed at 2 days post-inoculation (dpi) and 4 dpi; and (T5) pre-inoculation treatments, where hpTR-1+2 was sprayed 2 days and 4 days before virus infection **(Fig. 5).**

### Detection of MYMIV and Disease Severity Analysis

The progression of YMD symptoms in the inoculated plants was monitored on a daily basis. Disease severity scoring was performed according to a standardized 0-5 scale, as described in previous studies (59,64). Genomic DNA was isolated from the agroinoculated leaves (local) at 7 dpi and from the systemic leaves at 22 dpi. Rolling circle amplification (RCA) was conducted to amplify the MYMIV genomes. Subsequently, the RCA products containing MYMIV genomic DNA were monomerized through restriction digestion using PstI, a unique cutter. To detect the presence of the virus in the inoculated test plants, diagnostic conventional PCR analysis was performed using a specific primer set targeting the MYMIV DNA A region (**Table 2).**

**Table 2.**
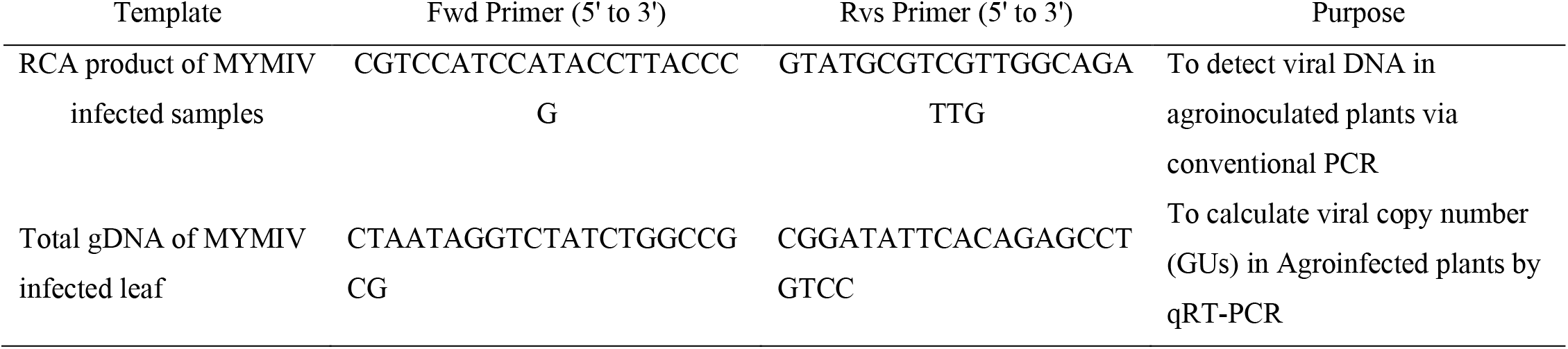
List of primers used to detect MYMIV genome and qRT-PCR.

To quantify the virus titre in the inoculated plants, an absolute quantification real-time PCR (qRT-PCR) experiment using the standard curve method was conducted. Standard curves were generated by utilizing plasmid containing the cloned full-length MYMIV DNA A genomic component, following the method described by (59,65,66). Tenfold serial dilutions of these plasmid, ranging from 10^2^ to 10^8^ copies of the viral genome per sample, were prepared in DNA extracted from non-infected mungbean plants and used as standards. Each qPCR reaction was performed in a 20 μl reaction volume, consisting of 10 μl of SYBR Green mix (PowerUp™ SYBR™ Green Master Mix, Applied Biosystems™), 1.6 μl of Rep (AC1) gene-specific primers **(Table 2),** and 1 μl of template DNA. The reaction conditions were optimized for primer pair, involving an initial denaturation at 95 °C for 2 minutes, followed by 40 cycles of denaturation at 95 °C for 15 seconds and annealing/extension at 60 °C for 30 seconds, using a Rotor-Gene® Q instrument (QIAGEN). The standard curve was generated by performing linear regression analysis of the Ct values plotted against the log of total DNA content in each dilution. To ensure accuracy and reproducibility, all reactions were performed with three technical and biological replicates.

Finally, viral gene expression analysis carried out using semi quantitative reverse transcriptase-PCR. Briefly, total RNA extracted from the systemic leaves (22 dpi) using Invitrogen™ TRIzol™ Reagent (Catalog number, 15596026). Then cDNA synthesis performed using Thermo Scientific™ RevertAid™ H Minus First Strand cDNA Synthesis Kit (Catalog number, #K1632) in a 20 μl reaction volume containing random hexamer primer. The reaction mixture was incubated for 5 min at 25°C followed by 60 min at 42°C. Next, PCR was performed using 2X G9 Taq PCR Master Mix (Catalog number, #G7117) with the (Rep) or AC1 gene specific primer to detect target viral gene expression and with the *Vigna radiata* tubuline gene (*vr*Tubulin) specific primer as an internal control.

### Statistical Analysis

The statistical analysis of the experimental data followed a rigorous methodology to ensure robust and reliable results. Mean values with standard error bars were utilized for clear data presentation. Statistical comparisons between mean values were conducted using both the Student T-test and ANOVA F-test, depending on the nature of the analysis. All statistical tests were conducted using Graph Prism 9.0 analytical software (La Jolla, CA). For qRT-PCR analysis, all treatment combinations were treated as independent experiments, and statistical analysis was performed by ANOVA, considering a *p*-value < 0.05 as indicative of significance.

## RESULTS

### Identification of Conserved Target Regions (TRs) in Begomoviruses

The study aimed to establish broad spectrum resistance against YMD in mungbean by identifying highly conserved genomic regions associated with MYMV and MYMIV. Analysis of these viruses involved in-depth examination of their complete genome sequences to identify potential targets for Host-Induced Gene Silencing (HIGS). Multiple sequence alignment analysis unveiled two conserved target regions, TR-1 (286 to 524 bp, orange color) and TR-2 (1141 to 1379 bp, green color), situated at distinct positions within the MYMIV DNA A genome **(Fig. 1).** TR-1 and TR-2 were selected based on their exceptional sequence conservation across various begomoviruses. TR-1 displayed a high sequence percent identity with MYMV (95 to 100%), HgYMV (∼88%), MYMIV (∼86 to 88%), TYLCV (∼84%), and Tomato leaf curl virus (TLCV, ∼81%) **(supplemental Table 1).** Similarly, TR-2 exhibited robust sequence conservation with MYMV (97-100%), MYMIV (90-98%), and HgYMV (91-94%) **(supplemental Table 1).**

**Fig. 1.**
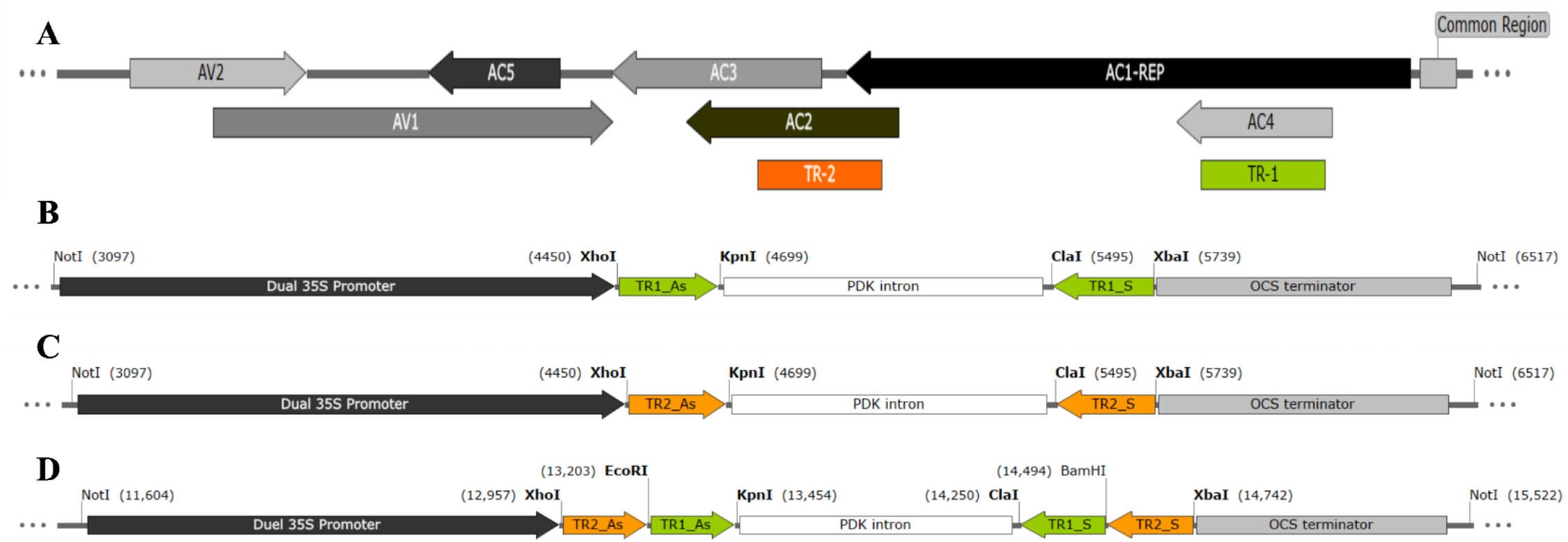
Schematic of MYMIV DNA A and three hpRNAi constructs. **A**) A complete MYMIV DNA A genome (GenBank Accessions: OK431083) showcasing various ORFs (AC1, AC2, AC3, AC4, AC5, AV1, and AV2) and the two selected target regions (TRs): TR-1 (AC4/AC1 genes; green color) and TR-2 (AC2/AC3/AC1 genes; orange color) position is shown. Schematic T-DNA map of pART27 RNAi cassettes of TR-1 (**B**), TR-2 (**C**), and TR-1+2 (**D**) in sense (S) and antisense (As) orientation. Abbreviations: Dual 35S Promoter: Cauliflower mosaic virus 35S promoter; OCS terminator: octopine synthase terminator, PDK intron: pyruvate dehydrogenase kinase intron, Restriction enzyme NotI were used for cloning of all the three RNAi cassettes from the intermediate RNAi vector pKANNIBAL into the plant transformation binary vector pART27.

### Differential Inhibition of MYMIV by Three hpRNAi Constructs in Mungbean

An evaluation was conducted to determine the efficacy of various constructs targeting MYMIV in the mungbean-MYMIV pathosystem. Response assessment and virus content monitoring were performed at 7 days post-inoculation (dpi) for inoculated leaves and at 22 dpi for systemic leaves, utilizing PCR and RCA-based restriction digestion analysis.

When mungbean plants were agroinoculated with MYMIV and hpTR-1 (AC4/AC1 hpRNAi construct), an average infection rate of 1 out of 10 was observed, without any visible symptoms **(Table 4).** The infection had a severity grade of 1, and though viral DNA was detected in local tissue at 7 dpi, it was notably absent in systemic leaves at 22 dpi. The qRT-PCR based quantification assays indicated a viral titre of less than 100 genomic units (GUs), indicating a highly resistant condition (**Table 3).** In another scenario involving MYMIV and hpTR-2 (AC2/AC3/AC1 hpRNAi construct) agroinoculation, the infection rate was 2 out of 10, accompanied by mild yellow mosaic symptoms with a severity grade of 2. Both PCR and RCA-based analyses confirmed viral presence in both local and systemic leaves, with a viral titre between 1000 and 8000 GUs, suggesting this construct’s ability to impart resistance against MYMIV. Utilizing hpTR-1+2 resulted in no observed infection among the 10 plants. No visible symptoms were recorded, and both PCR and RCA analyses displayed the absence of viral DNA in local and systemic leaves at 7 and 22 dpi, with negligible viral titre **(Fig. 2).** Notably, it was found that hpTR-1+2 effectively restricts the systemic movement of MYMIV and its viral gene expression **(Fig. 3 C).** This illustrated the robust immunity conferred by hpTR-1+2_pART27 (AC4/AC1**+**AC2/AC3/AC1) stack construct. Conversely, agroinoculation with MYMIV and an empty vector led to infections in all plants, exhibiting severe YMD symptoms with a severity grade of 5 **(Table 4).** Both PCR and RCA analyses revealed the presence of viral DNA in both local and systemic leaves, with a viral titre surpassing 30000 GUs, indicating high susceptibility.

**Fig. 2.**
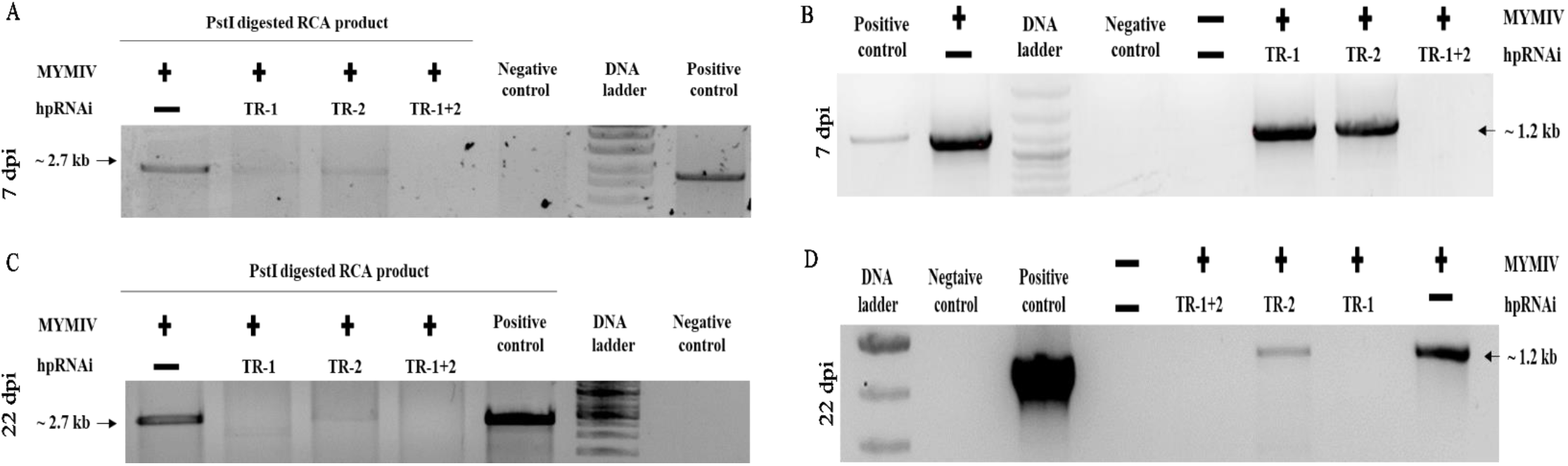
Resistance imparted by three hpRNAi constructs against MYMIV. RCA analysis: RCA product generated from total genomic DNA of mungbean leaf collected at local leaves at 7 dpi (**A**) and systemic leaves at 22 dpi (**C**) from hpRNAi transient assay. RCA product subjected to restriction digestion using a unique cutter (pstI). Appearance of a 2.7 kb fragment indicates the presence of MYMIV DNA A components; Conventional PCR analysis: Detection of viral DNA using a MYMIV DNA-A specific primer. Genomic DNA extracted from mungbean leaf collected at local leaves at 7 dpi (**B**) and systemic leaves at 22 dpi (**D**) from hpRNAi transient assay.

**Fig. 3.**
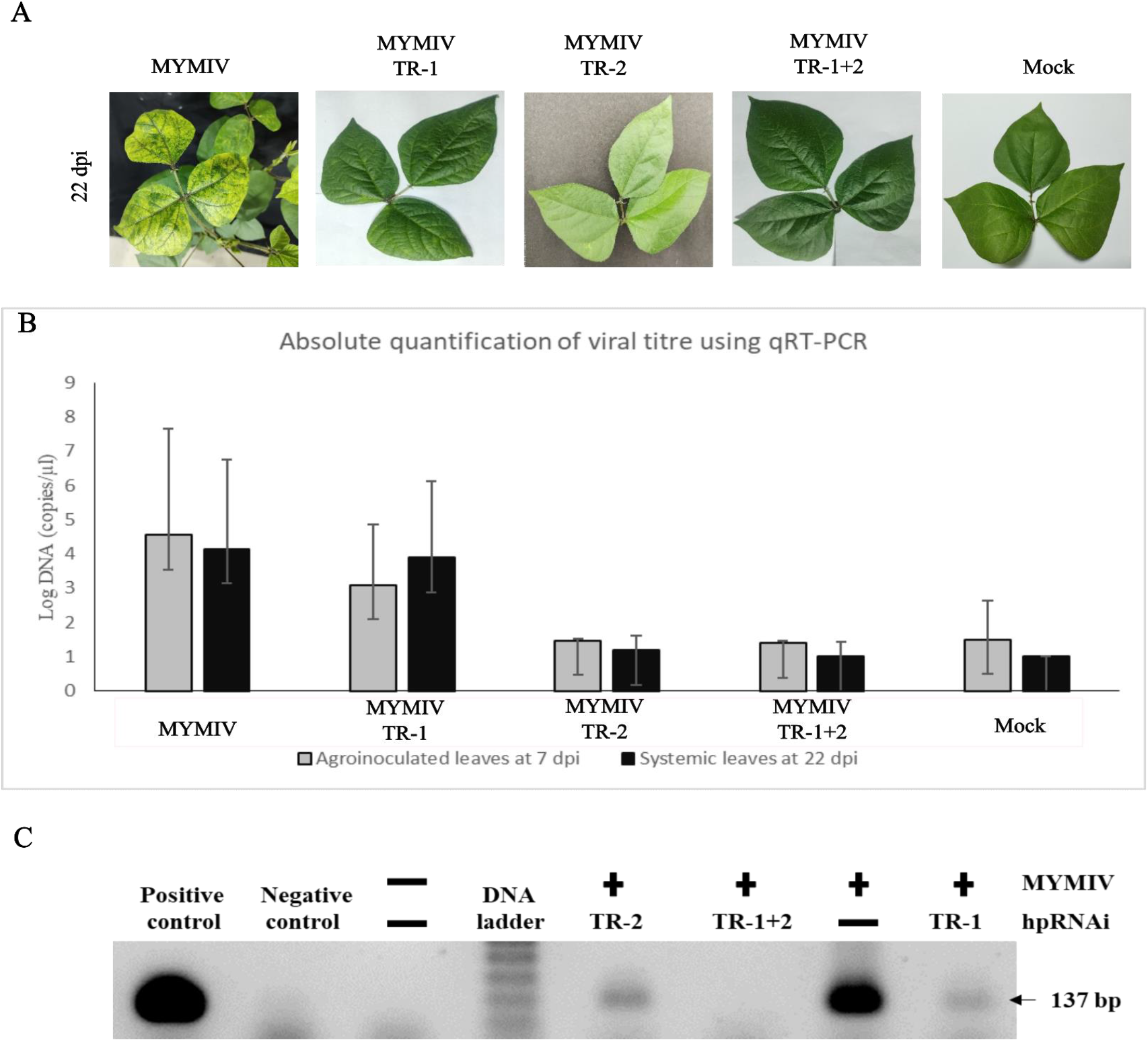
Efficacy of three hpRNAi constructs against MYMIV infection in mungbean. **A**) Systemic symptom analysis: Evaluation at 22 dpi on mungbean plants treated with hpTR-1, hpTR-2, and hpTR-1+2 transient expression and agroinoculated with MYMIV; **B**) Viral titre measurement by qRT-PCR: Quantification in local leaves at 7 dpi and systemic leaves at 22 dpi. Data points represent means of triplicate measurements, and vertical lines on each bar indicate standard deviations. Statistically significant difference indicates (*p* < 0.05, ANOVA); **C**) Viral gene expression analysis by semi-quantitative RT-PCR: Utilizing AC1 gene-specific primer. Detection of a 137 bp fragment in systemic leaves at 22 dpi indicates the presence of viral mRNA.

**Table 3.**
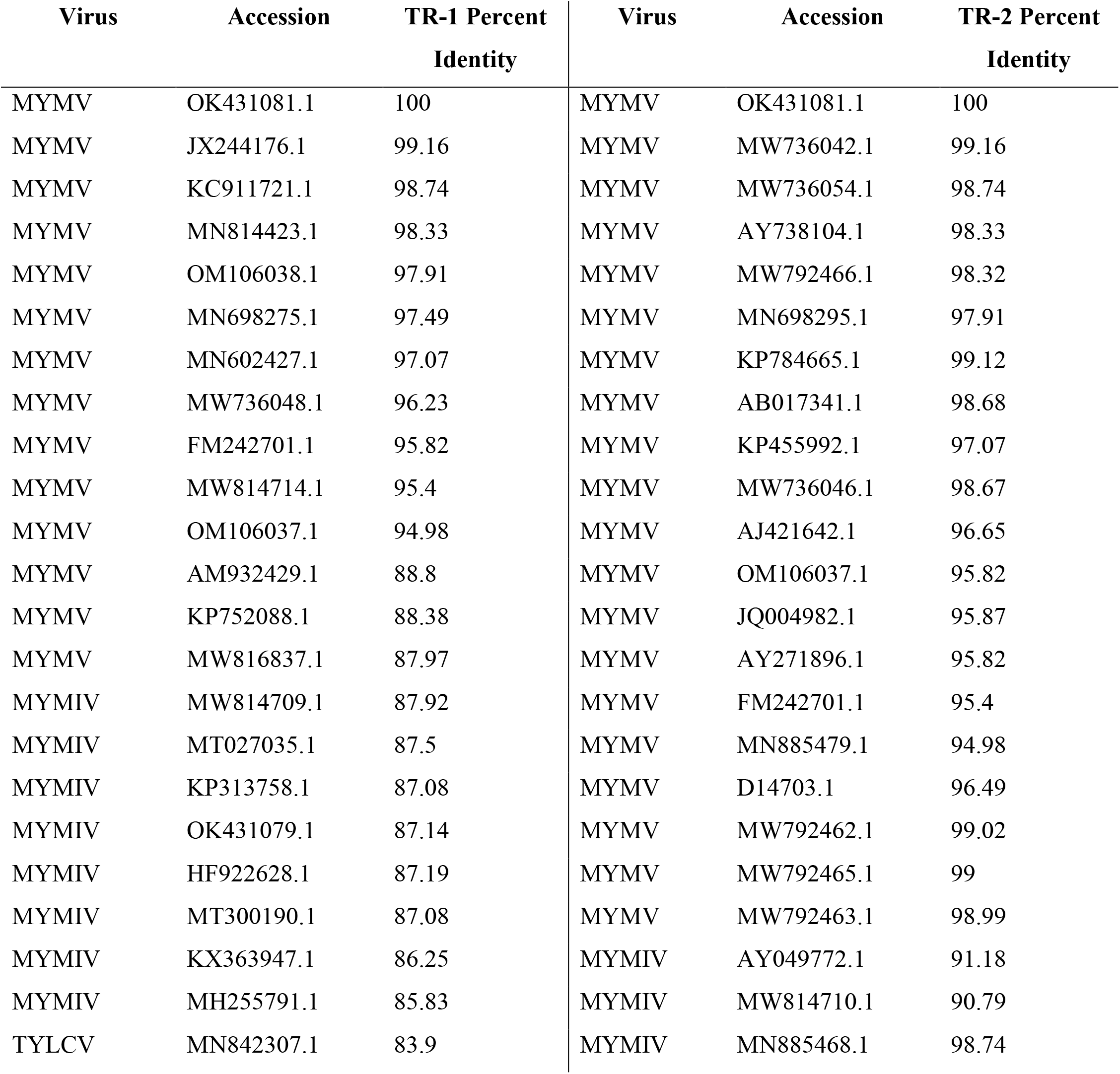
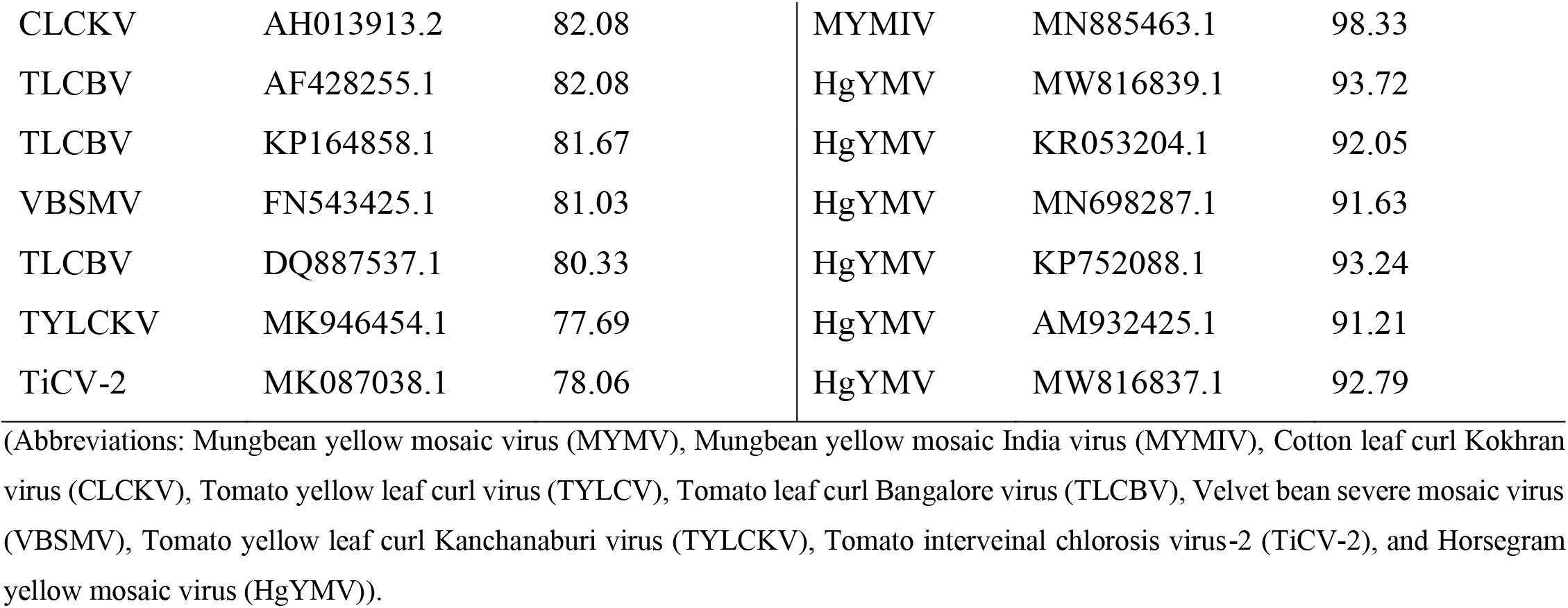
TR-1 and TR-2 sequence percent Identity. The percent sequence identity analysis of selected RNAi target regions with begomoviruses from various plant using NCBI BLASTn.

**Table 4.**
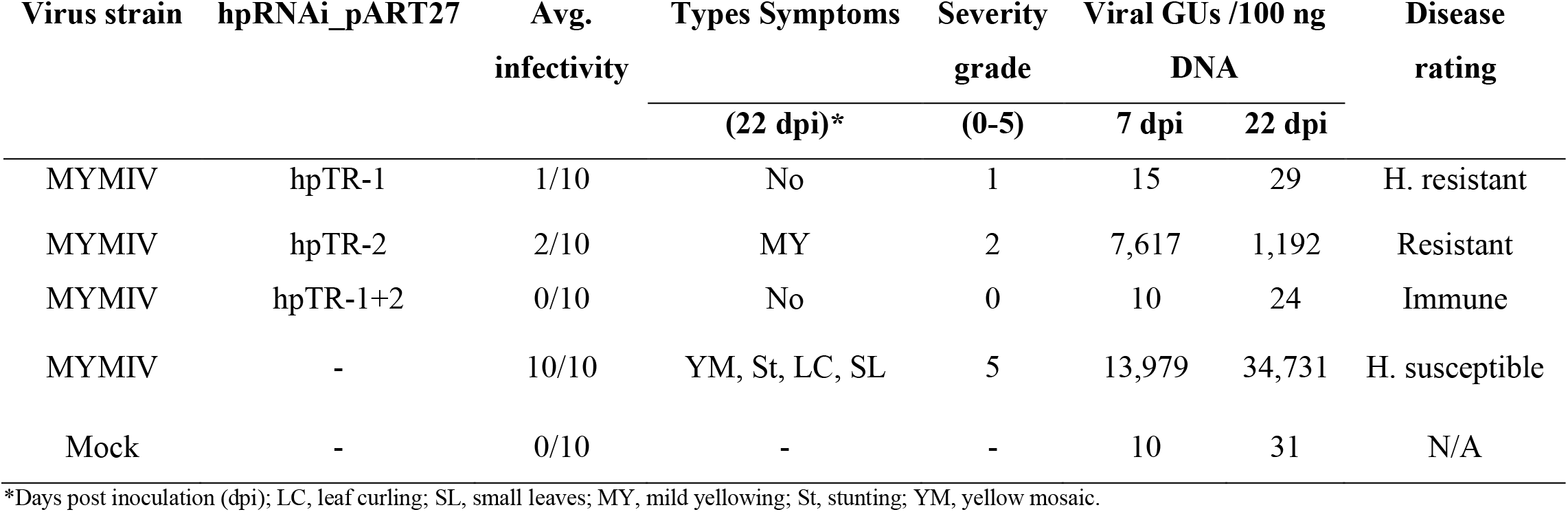
Infectivity analysis of MYMIV in hpRNAi transient expression assay in mungbean.

| Virus strain | hpRNAi_pART27 | Avg.<br>infectivity | Types Symptoms<br>(22 dpi)* | Severity<br>grade<br>(0-5) | Viral GUs /100 ng<br>DNA |  | Disease<br>rating |
| --- | --- | --- | --- | --- | --- | --- | --- |
|  |  |  |  |  | 7 dpi | 22 dpi |  |
| MYMIV | hpTR-1 | 1/10 | No | 1 | 15 | 29 | H. resistant |
| MYMIV | hpTR-2 | 2/10 | MY | 2 | 7,617 | 1,192 | Resistant |
| MYMIV | hpTR-1+2 | 0/10 | No | 0 | 10 | 24 | Immune |
| MYMIV | - | 10/10 | YM, St, LC, SL | 5 | 13,979 | 34,731 | H. susceptible |
| Mock | - | 0/10 | - | - | 10 | 31 | N/A |

\*Days post inoculation (dpi); LC, leaf curling; SL, small leaves; MY, mild yellowing; St, stunting; YM, yellow mosaic.

### Selection and Synthesis of hpRNA

The mungbean-MYMIV pathosystem, coupled with Agrobacterium-mediated transient expression assays, pinpointed the highly effective hpTR-1+2 construct. This transient assay revealed varied inhibition levels of MYMIV in mungbean leaves, with the hpTR-1+2 clone demonstrating superior effectiveness in impeding MYMIV accumulation and systemic movement compared to individual hpTR-1 and hpTR-2 constructs. Consequently, the hpTR-1+2 cassette was selected for hpRNA production. The cassette was obtained as an XhoI and XbaI fragment from the hpTR-1+2_pART27 clone and subcloned into the L4440 vector. The resulting plasmid, hpTR-1+2_L4440, was then transformed into *E. coli* strain HT115 for *in vivo* expression **(Fig. 4 A)**. In the *in vivo* experiment, 1 to 1.5 mg of hpRNA was obtained from a 50 ml bacterial culture after RNase A digestion. The quality of hpTR-1+2 was assessed on a 1.5% agarose gel, and its concentration was measured using a nanodrop. Purified hpTR-1+2 underwent nuclease treatments to confirm its double-stranded nature **(Fig. 4 B).** Finally, sequence confirmation was achieved through semi-qRT-PCR analysis, using the AC2-S2-XhoI-FP and AC4-S2-kpnI-RP primer pair, amplifying a 500 bp fragment of the sense strand of hpTR-1+2 **(Fig. 4 A and B).**

**Fig. 4.**
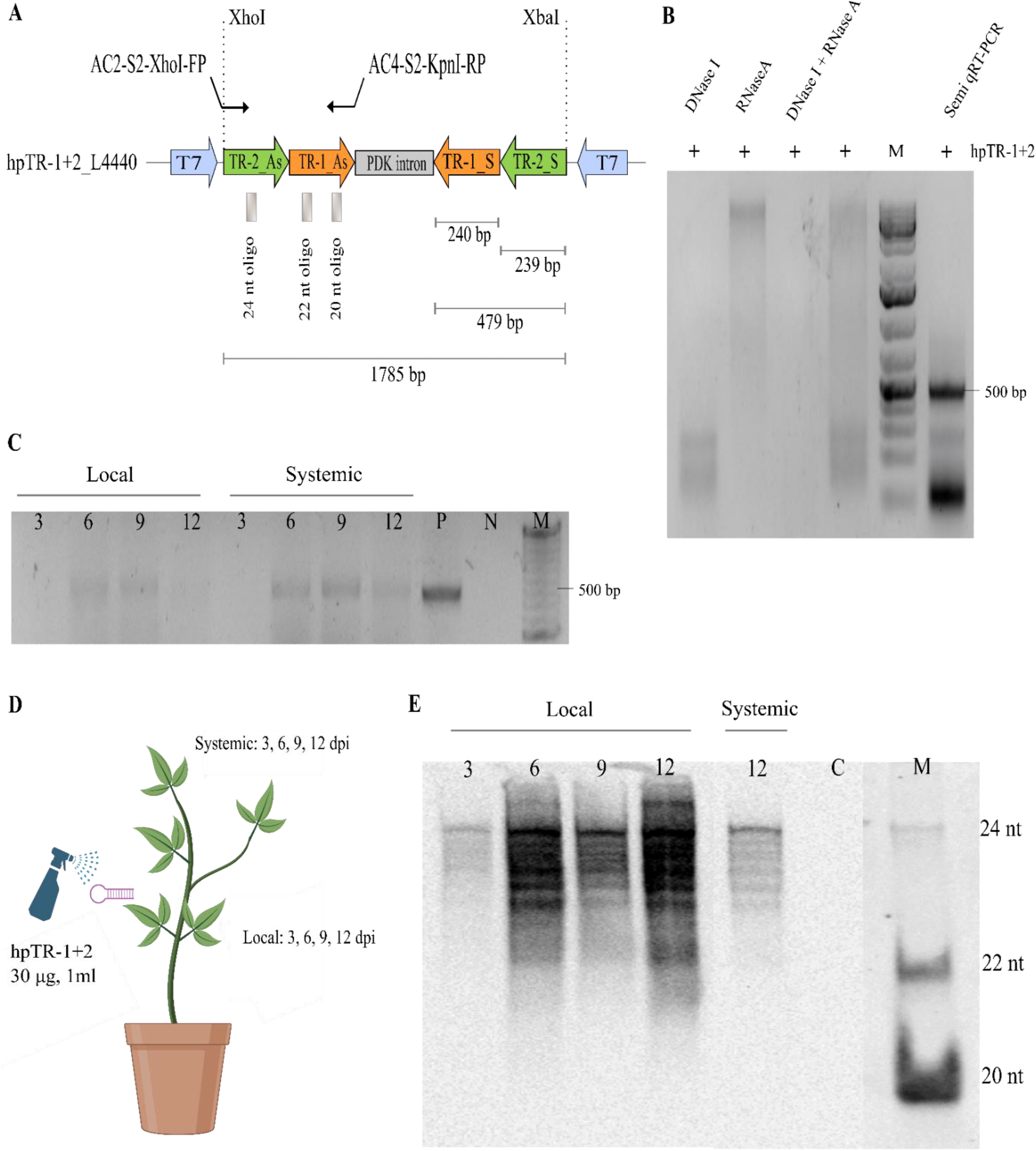
In vivo production of hpRNA, persistence, systemic movement, and induction of siRNA formation. **A)** Schematic representation illustrates hpTR-1+2_L4440 plasmid: The highly efficient hpRNA cassette (hpTR-1+2) is subcloned into the L4440 plasmid, a vector with two T7 promoters. This plasmid construction is achieved through the utilization of XhoI and XbaI restriction sites. The resulting plasmid, designated hpTR-1+2_L4440, is subsequently transformed into *E. coli* strain HT115 to facilitate *in vivo* expression; **B)** *In vivo* production of hpRNA in bacteria: In this step, the *E. coli* strain HT115, hosting the hpTR-1+2_L4440 construct, is induced with 0.4 mM IPTG to initiate the *in vivo* expression of hpTR-1+2. Post-induction, total RNA was isolated and subjected to DNase I and RNase A treatment to confirm the double-stranded nature of the RNA. The hpRNA sequence was verified through semi-quantitative reverse transcriptase PCR (semi-qRT-PCR) using the AC2-S2-XhoI-FP and AC4-S2-kpnI-RP primer pair, amplifying a 500 bp fragment of the sense strand of hpTR-1+2. The resulting hpRNAs were electrophoresed through a 1.5% agarose gel and visualized by staining with ethidium bromide. Lane M represents 1 kb Plus DNA Ladder (Invitrogen); **C)** Detection of hpTR1+2 in local (treated) and systemic (non-treated) leaves of mungbean plants by semi-qRT-PCR analysis: This phase involves the validation of hpRNA internalization into plant cells and the assessment of its persistence and systemic movement. Leaf samples obtained from both local and systemic regions undergo semi-qRT-PCR analysis. Positive (P) and negative (N) controls are included for reference, and a DNA ladder (M); **D)** Treatment of mungbean plants and sample collection: A comprehensive schematic diagram outlines the treatment procedure of three-week-old mungbean plants. These plants are sprayed with a solution containing 30 µg/ml of hpRNA on leaves without MYMIV infection. Subsequent leaf sample collected at various time points, spanning 3, 6, 9, and 12 days-post-spray, from both local and systemic trifoliate. These samples are destined for further semi-qRT-PCR and northern blot analyses; **E)** Detection of hpTR1+2-derived small interfering RNAs (siRNAs) in local and systemic leaves of mungbean plants by northern blot analysis: This final step involves assessing the ability of hpRNA to initiate RNA interference (RNAi) in plant cells, leading to the induction of small interfering RNA (siRNA) formation. The northern blot analysis meticulously examines siRNA size, persistence, and systemic movement. Lane C represents non-treated control plants, while Lane M consists of three oligomers (each with lengths of 24 nt, 22 nt, and 20 nt), serving as siRNA size references.

### Cellular Uptake, Systemic Movement, and siRNA Induction by hpRNA

Hairpin RNA (hpTR-1+2) was externally applied to trifoliates of 2-3-week-old mungbean plants without MYMIV infection to explore its cellular uptake, stability, and systemic movement. Purified naked hpTR-1+2 at a concentration of 30 µg/ml was sprayed onto the mungbean plants (**Fig. 4 D**). The abundance of hpRNA was assessed at various time points (3, 6, 9, and 12 days-post-spray) from both local (treated) and systemic (non-treated) trifoliates using semi-quantitative reverse transcription-polymerase chain reaction (RT-PCR) analysis (performed same as above). The results demonstrated the presence of hpTR-1+2 in both local and systemic tissues up to 12 days post-spray (**Fig. 4 C)**. This data indicates that the externally applied hpRNA is effectively taken up by plant cells, maintains stability for up to 12 days, and exhibits systemic movement from the application site to fresh leaves.

To assess the potential of exogenously applied hpRNA molecules to generate siRNAs in planta, their detection was attempted through northern blot analysis. Its presence was monitored in treated (local) leaves at 3, 6, 9, and 12 days-post-spray, as well as in non-treated (systemic) leaves at 12 days post-spray in mungbean. Northern blot analysis utilized both sense and antisense strands of hpTR-1+2 as a DIG-labeled hybridization probe for siRNA detection. The northern blot data suggested the presence of siRNAs ranging from 24 to 22 nucleotides in length, with the intensity of siRNA signals markedly increasing from 6 days post-spray in local tissues **(Fig. 4 E**). In systemic tissues at 12 days post-spray, a lower intensity of a similar siRNA pattern was observed. This data strongly indicates that exogenously applied hpRNA effectively initiates RNAi, leading to the generation of siRNAs in planta, and induces their formation in both local and systemic tissues.

### Efficacy of Spray-Induced Gene Silencing Against MYMIV in Mungbean

To optimize the efficacy of hpRNA-based vaccination, five treatment combinations (refer to **Table 5**) were employed with MYMIV inoculation. Disease incidence was assessed at systemic leaves 24 days post-MYMIV inoculation using PCR and RCA-based restriction digestion analysis.

**Table 5.**
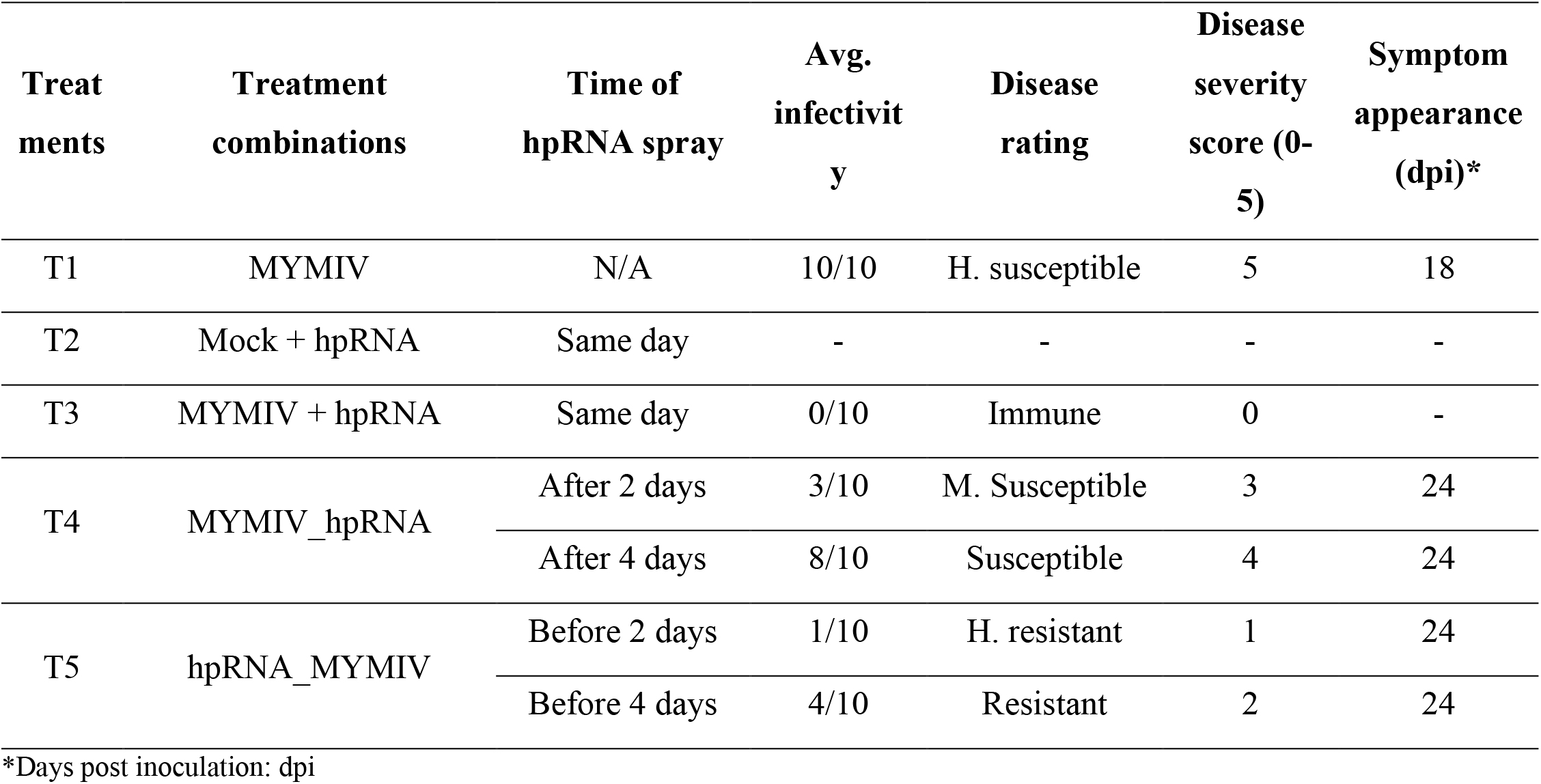
Overview of hpRNA spray treatment combinations and YMD incidence.

| **Treatments** | **Treatment combinations** | **Time of hpRNA spray** | **Avg. infectivity** | **Disease rating** | **Disease severity score (0-5)** | **Symptom appearance (dpi)\*** |
| --- | --- | --- | --- | --- | --- | --- |
| T1 | MYMIV | N/A | 10/10 | H. susceptible | 5 | 18 |
| T2 | Mock + hpRNA | Same day | - | - | - | - |
| T3 | MYMIV + hpRNA | Same day | 0/10 | Immune | 0 | - |
| T4 | MYMIV_hpRNA | After 2 days | 3/10 | M. Susceptible | 3 | 24 |
|  |  | After 4 days | 8/10 | Susceptible | 4 | 24 |
| T5 | hpRNA_MYMIV | Before 2 days | 1/10 | H. resistant | 1 | 24 |
|  |  | Before 4 days | 4/10 | Resistant | 2 | 24 |

\*Days post inoculation: dpi

Mungbean plants inoculated with MYMIV without hpRNA spray exhibited typical YMD symptoms at 18 dpi, consistent with the expected timeframe for symptom appearance (**Fig. 5 A**). All plants in this treatment displayed 100% disease incidence, as confirmed by conventional PCR analysis, revealing the anticipated 1.2 kb fragment (**Fig. 5 B**). Additionally, RCA-based restriction digestion identified a characteristic 2.7 kb band (**Fig. 5 C**), affirming the infectious nature of the MYMIV inoculum and suitable environmental conditions for symptom development on mungbean plants. In contrast, mock-inoculated plants treated with hpRNA spray, using an empty pCambia3300 vector instead of the vector carrying the infectious dimeric clone of MYMIV, remained healthy without exhibiting any symptoms.

**Fig. 5.**
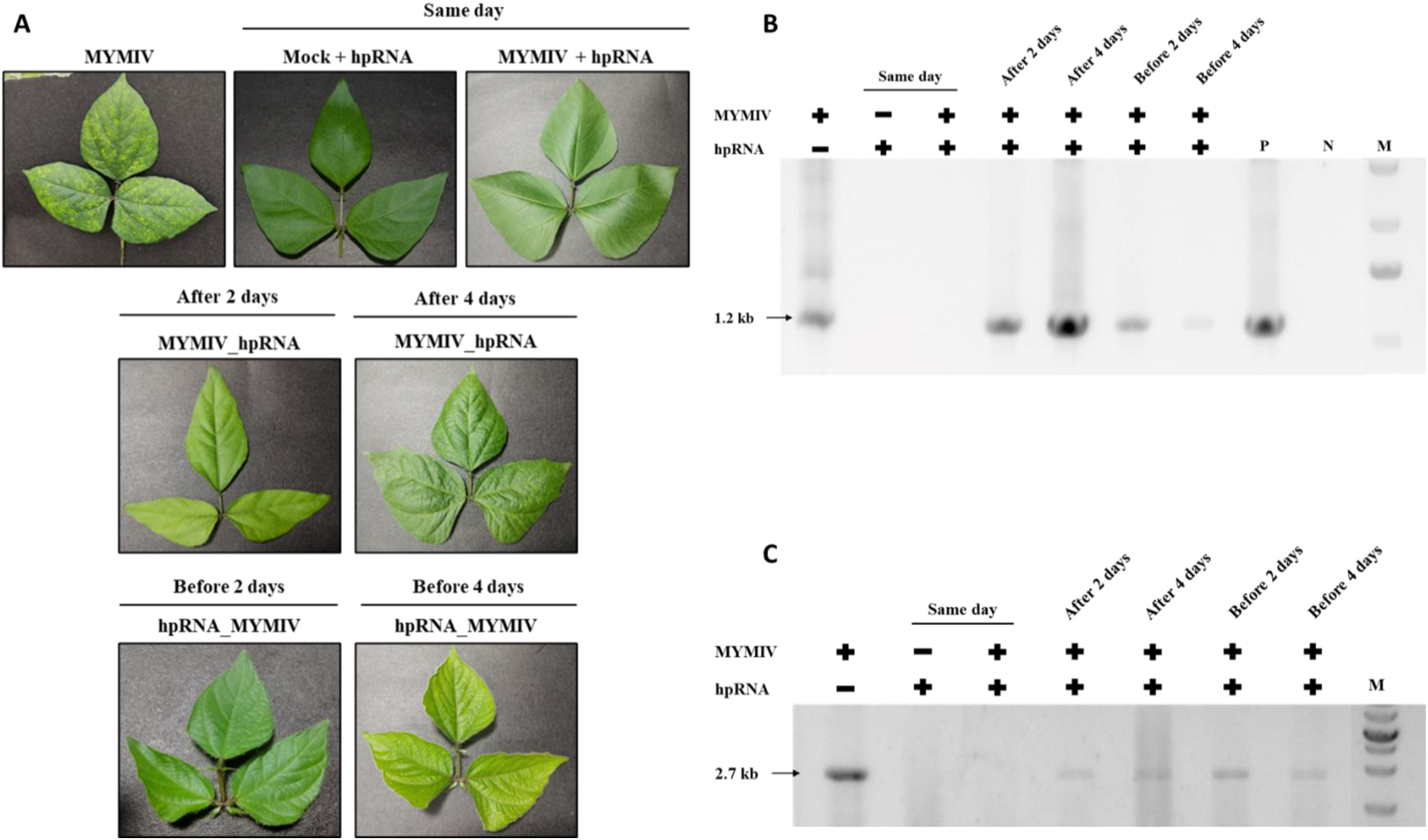
In vivo-produced hpRNA confers resistance to MYMIV in mungbean. **A)** Phenotypic response of mungbean plants to MYMIV agroinoculation with in vivo-produced hpRNA. Inoculation scenarios: MYMIV agroinoculation without hpRNA (positive control); mock inoculation followed by 30 µg hpTR-1+2 spray; MYMIV infection, with immediate hpTR-1+2 spray; topical hpTR-1+2 spray at 2 and 4 dpi after MYMIV agroinoculation; pre-inoculation hpTR-1+2 spray 2 and 4 days before virus infection. Systemic symptoms were photographed at 24 dpi; **B)** Conventional PCR analysis: Detection of viral DNA using MYMIV DNA-A specific primer. Genomic DNA extracted from mungbean leaf collected at systemic leaves at 24 dpi from hpRNA spray assay. N, negative control (water); P, positive control (pCAMBIA 3300 harboring MYMIV DNA A); and M, DNA Ladder; **C)** RCA analysis: RCA product generated from total genomic DNA of mungbean leaf collected at systemic leaves at 24 dpi from hpRNA spray assay. RCA product subjected to restriction digestion using a unique cutter (PstI). Appearance of a 2.7 kb fragment indicates the presence of MYMIV DNA A components.

Despite the topical application of single hpRNA molecules with sequence homology to key genes of MYMIV, variable levels of protection against MYMIV infection were observed in three repeated experiments. The co-application of in vivo produced hpTR-1+2 together with MYMIV on plants provided 100% protection, as evident by absence of DNA fragments in PCR and RCA based analysis and complete lack of YMD symptoms. Notably, this data suggests that spraying hpRNA on the same day as infection provides highest protection (**Table 5**).

The protective efficacy of hpRNA was further assessed by topical spray of hpRNA after MYMIV agroinoculation, performed at 2 dpi and 4 dpi. In this treatment, at 2 dpi, 30% of plants exhibited a disease severity of scale 3 out of 5, indicating moderate susceptibility to MYMIV. Conversely, when hpRNA was sprayed after MYMIV inoculation at 4 dpi, even less protection activity was observed. Nevertheless, hpRNA spray delayed the onset of YMD symptoms by six days, and viral DNA was detected by PCR and RCA-based analysis in systemic leaves at 24 dpi instead of 18 dpi.

The hpRNA-based vaccination activity of hpTR-1+2 was validated by topical spray of hpRNA before MYMIV agroinoculation, administered at 2 and 4 days before virus infection. Interestingly, spraying hpRNA two days before virus infection increased plants ability to significantly restrict viral accumulation and decreased disease severity by many fold in this treatment, only 10% of the plants were infected, exhibiting high resistance to MYMIV., viral DNA was detected by molecular analysis after delayed symptom appearance at 24 dpi. More interestingly 4 days before hpRNA spray also provided help combat virus infection showed MYMIV resistance with an average infectivity of 40% and disease severity of scale 2 out of 5. Viral DNA was detected by molecular analysis after delayed symptom appearance at 24 dpi. From these results, it can be inferred that the hpRNA spray treatment before virus infection provided a certain level of protection against MYMIV infection in mungbean (**Table 5**).

Intriguingly, spraying hpRNA two days before virus infection significantly enhanced the plant’s ability to restrict viral accumulation and decreased disease severity by many folds. In this treatment, only 10% of the plants were infected, exhibiting high resistance to MYMIV, with viral DNA detected after a delayed symptom appearance at 24 dpi. Similarly, spraying hpRNA 4 days before virus infection provided moderate protection, resulting in an average infectivity of 40% and a disease severity of scale 2 out of 5. Viral DNA was detected after delayed symptom appearance at 24 dpi. These results collectively suggest that hpRNA spray treatment before virus infection confers a certain level of protection against MYMIV infection in mungbean.

## DISCUSSION

RNA interference, triggered by dsRNA molecules, is a potent method for controlling plant viruses in transgenic plants (67). However, global approval of transgenesis is lacking, and concerns about potential adverse effects on the natural environment exist. As an alternative, a non-transgenic approach utilizing RNAi, known as ‘RNA-based vaccination,’ has been developed for the control of plant viruses (46). Tenllado and Díaz-Ruíz, 2001 were pioneers in reporting antiviral protection through the topical application of homologous dsRNA (46). Since then, various studies have reported the effectiveness of directly applying dsRNA to confer resistance against a broad spectrum of RNA viruses (46,49,51,53,54,68–72). In the context of DNA viruses, the first study came from Namgial et al. (2019), they reported protection against a bipartite geminivirus, ToLCV in tomato plants through the direct application of dsRNAs. Their study demonstrated significant protective effects, with 45%, 60%, and 50% reductions in disease incidence when dsRNAs targeting AC1/AC4, AV1/AV2, and AC1/AC4_AV1/AV2, respectively, were applied directly (54). On the other hand, Melita et al. 2021 provided the first report on monopartite geminivirus protection. In their study, dsRNAs were synthesized in vitro, targeting the C4 and V2 genes of TYLCV. When these dsRNA molecules were topically applied onto tomato plants along with the virus (via agroinfiltration), they observed a significant reduction in disease incidence to 23% and 46%, respectively, against TYLCV (56). Spray application of a in vivo produced dsRNA cocktail targeting suppressor genes C2, V2, and C4 reduced ChiLCV (a monopartite begomovirus) incidence by up to 66.7% in *Nicotiana benthamiana* (57). In a recent study (Krishnamoorthy et al., 2023), in vitro synthesized dsRNA targeting the coat protein gene (CP) and replication initiator protein gene (Rep) of MYMV was exogenously applied (2.5 μg of dsRNA) to blackgram plants showing MYMV symptoms. This treatment led to a reduction in disease severity for several days. Among plants treated with dsRep, 80% exhibited symptom remission at the 21st day post dsRNA treatment, while for dsCP treatment, 73% of plants showed symptom remission at the 15th day (55). Though, exogenously applied dsRNA presents a promising tool for virus control, its efficacy may vary across different viruses. For DNA viruses like the begomoviruses Tomato Severe Rugose Virus (ToSRV) in tomatoes (73) and Tomato Leaf Curl New Delhi Virus (ToLCNDV) in zucchini squash (50), the application of exogenous dsRNA has shown limited effectiveness. Considering the existing research gap, this study aimed to assess the effectiveness of exogenously applied hpRNA for the control of a bipartite begomovirus, MYMIV, in economically significant mungbean plants.

In this study, two RNAi target regions for HIGS in the MYMIV genome were identified: TR-1, targeting AC4/AC1 genes, and TR-2, targeting AC3/AC2/AC1 genes (Fig. 1 A). In most studies, these regions were selected due to their high conservation within the viral genome **(Table 3**), as demonstrated in previous studies that achieved effective resistance against begomoviruses through both transgenic and transient HIGS approaches (41,74–80). Three hairpin RNA interference (hpRNAi) constructs were prepared using these two target regions to evaluate their efficacy against MYMIV in a transient HIGS assay (**Fig. 1 B, C, and D)**. The data indicated that the hpTR-1+2 clone (a stacked clone of AC4/AC1_AC3/AC2/AC1) was the most effective, providing immunity against MYMIV in mungbean compared to individual clones **(Fig. 2), (Fig. 3),** and **(Table 4).** Consequently, the hpTR-1+2 clone was selected for in vivo dsRNA synthesis for the spray induced gene silencing (SIGS) assay.

In the process of inducing RNAi through topical application, the preparation of dsRNA is a crucial step. The dsRNA can be efficiently produced in vivo using bacterial expression systems (81,82), which offer convenience and economic advantages compared to in vitro methods involving RNA polymerase (83). Here, we employed a *E. coli* HT115 expression system to generate hpRNA (hpTR-1+2) in large quantities. The dose of exogenous dsRNA varied across studies: 40 to 60 μg against Zucchini Yellow Mosaic Virus (ZYMV) (51), almost 200 μg/plant against Tobacco Mosaic Virus (TMV) (49), 2.5 μg/plant against MYMV (55), 250 μg/plant against Tomato Spotted Wilt Virus (TSWV) (72), 60 μg/plant against Cucumber Green Mottle Mosaic Virus (CGMMV) (50), 15 μg/plant against ChiLCV (57), 100 μg/plant against Pepper Mild Mottle Virus (PepMoV) (48), 12 to 16 μg/plant against Cucumber Mosaic Virus (CMV) (53), 10 μg/plant against TSWV (70), and 20 to 30 μg/plant against ToLCV (54). Machado et al. (2020) found a threshold, with concentrations below 16 μg/plant providing no protection in Tobacco Mosaic Virus (ToMV)-dsRNA tests, and a dose-dependent response ranging from 50 to 400 μg/plant (52). Given this context, we utilized 30 μg/plant of hpRNA against MYMIV in mungbean **(Fig. 4 D**).

To optimize the effectiveness of hpRNA in suppressing MYMIV multiplication, the timing of virus and dsRNA application is crucial. In this study, mungbean cv. K851 was treated with the virus and hpRNA separately to determine the optimum treatment scenario. The most significant inhibition of MYMIV replication occurred when plants were treated with hpRNA on the same day, two days, and four days before viral inoculation, resulting in disease ratings of immune, highly resistant, and resistant, respectively **(Fig. 5)** and (**Table 5).** These treatments demonstrated higher protection efficiency compared to post-treating plants at two or four days after virus inoculation. The earlier introduction of hpRNA proved more effective, suggesting that prior treatment allows for the generation of siRNAs and the formation of RISC in advance, facilitating an immediate response upon virus infection. This underscores the importance of a certain period for hpRNA processing within cells to observe the RNAi effect of exogenous hpRNA treatment. This observation is consistent with findings from previous studies on other viruses treated with dsRNA (48).

The uptake mechanisms of exogenously applied dsRNAs remain partially understood, with absorption capacity varying among plant organs. Overcoming physical barriers like the wax layer, cuticle, cell wall, and cell membrane, successful external application of dsRNA in plants for inducing RNAi involves methods such as mechanical rubbing, pressure spray, infiltration, injection, root or petiole absorption, and nano-carrier conjugation (84). Here, leaf spraying was chosen as the method for its recognized effectiveness in inducing RNAi (85). The cellular mechanism of dsRNA-induced RNA interference (RNAi) in plants involves several steps: Cellular uptake of dsRNAs, followed by rapid cleavage by DICER-LIKE (DCL) endonucleases into 20 to 25-nucleotide siRNAs with 2-nt 3′ overhangs at both ends. Subsequently, one strand of siRNAs is incorporated into an ARGONAUTE (AGO) protein to form an RNA-induced silencing complex (RISC). Finally, the siRNA molecules guide the RISC to scan the cytoplasm for recognition and cleavage/degradation of complementary transcripts, leading to post-transcriptional gene silencing (PTGS) (86,87).

Despite the transient nature of protection, both hpRNA and siRNA molecules (21 to 25 nucleotides long) remained detectable on local and systemic leaves for at least 12 days, as shown in (**Fig. 4 C and E**). This observation suggests that a significant portion of hpRNA may have entered the leaf apoplast through spray application as it is the first barrier before entering inside cells, facilitating systemic transport to other parts of the plants. Microscopic analyses have demonstrated the presence of spray-applied dsRNAs in various plant structures, including the apoplast (88). Subsequently, this transport might have triggered RNAi in receiving plant cells, leading to the formation of siRNA from hpRNA. Our results indicate the need for a gradual and continuous introduction of dsRNA/hpRNA into the cells or the implementation of a sustained amplification mechanism for specific RNA processing machinery to achieve durable and efficient resistance in plants.

In this study, a short-term protection window was achieved with the use of naked hpRNAs spray approach. This limitation could be addressed by exploring alternative delivery methods, such as high-pressure spraying, and investigating nanoparticle-based delivery approaches. These methods have the potential to enhance the stability and efficacy of exogenously applied dsRNAs when compared to naked dsRNA delivery (89,90).

## CONCLUSION

In summary, our study marks the pioneering exploration of utilizing exogenous hairpin RNA (hpRNA) for managing yellow mosaic disease in mungbean induced by Mungbean Yellow Mosaic India Virus (MYMIV). By designing three hpRNAi constructs targeting key regions (TR-1: AC4/AC1, TR-2: AC3/AC2/AC1), we identified the most efficient clone through transient Host-Induced Gene Silencing (HIGS) assays—hpTR-1+2_pART27 (AC4/AC1_AC3/AC2/AC1). Notably, our study reveals a protection window with the naked hpRNA spray approach, particularly with in vivo-synthesized hpRNA (hpTR-1+2), showcasing its significant inhibition of MYMIV replication when applied on the same day, two days, or four days prior to viral inoculation. These findings bear considerable implications for the effective management of MYMIV in mungbean plants.

## Statements & Declarations

## Authors’ contributions

LS and KVD designed the research; KVD carried out the all experiments, analysed the data, and prepared the complete manuscript; LS corrected the manuscript. All authors read and approved the final manuscript.

## Acknowledgements

We are grateful to CSIRO, Australia for the pKANNIBAL and pART27 vectors. We express our sincere thanks to Prof. Sampa Das, Bose Institute for providing *Agrobacterium tumefaciens* strain, EHA105 and Center for Application of Molecular Biology to International Agriculture (CAMBIA), Australia for pCAMBIA3300. We also thankfull to Mr. Bharatheeswaran Murugan for his technical assistance.

## Availability of data and materials

The datasets used and/or analyzed during the current study are available from the corresponding author on reasonable request.

