## supplemental for "Hairpin-RNA Spray Confers Resistance to Mungbean Yellow Mosaic India Virus in Mungbean"

+91361-2582239 (Lab)

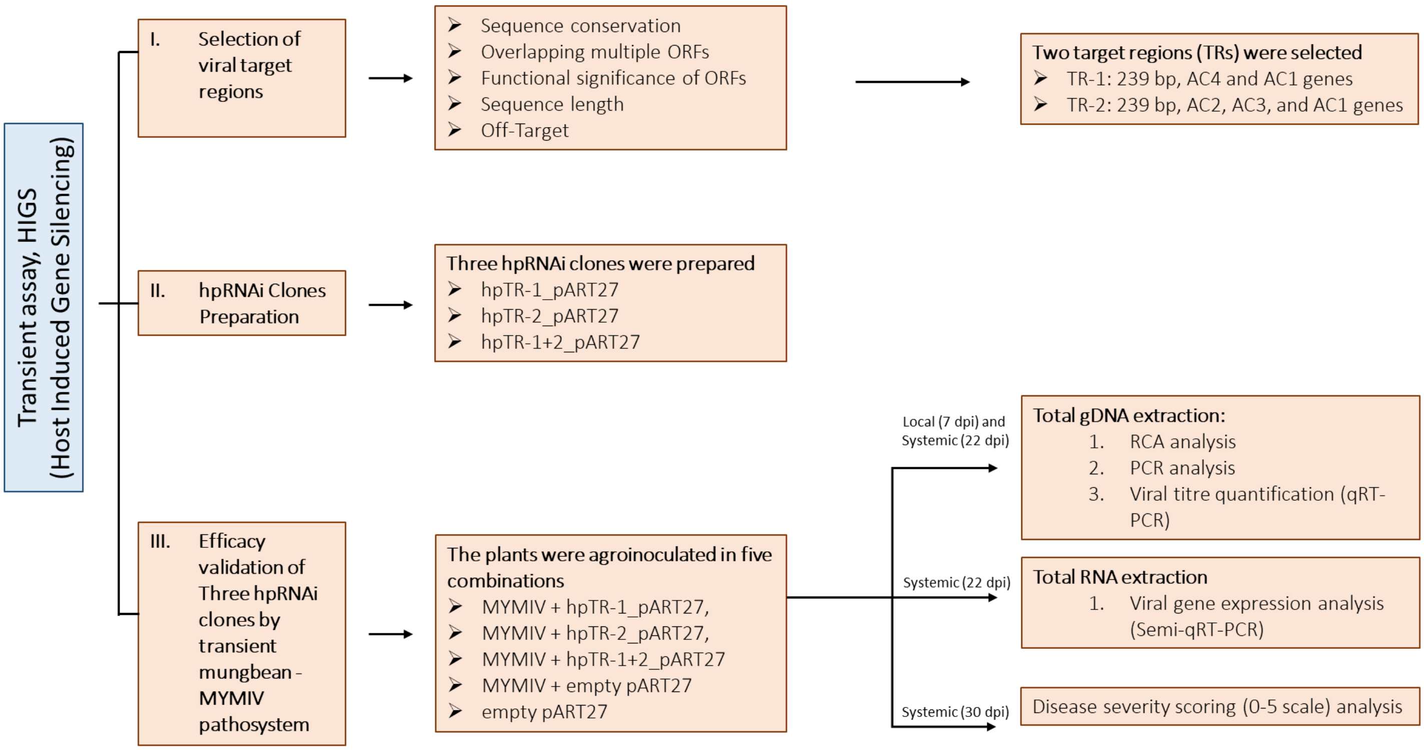


**Fig. S1: Schematic representation of the methodology adopted for the efficacy validation of three hairpin RNA interference (hpRNAi) constructs through transient expression in mungbean.**

The diagram outlines the key steps and components involved in the experimental approach to assess the effectiveness of the hpRNAi constructs.

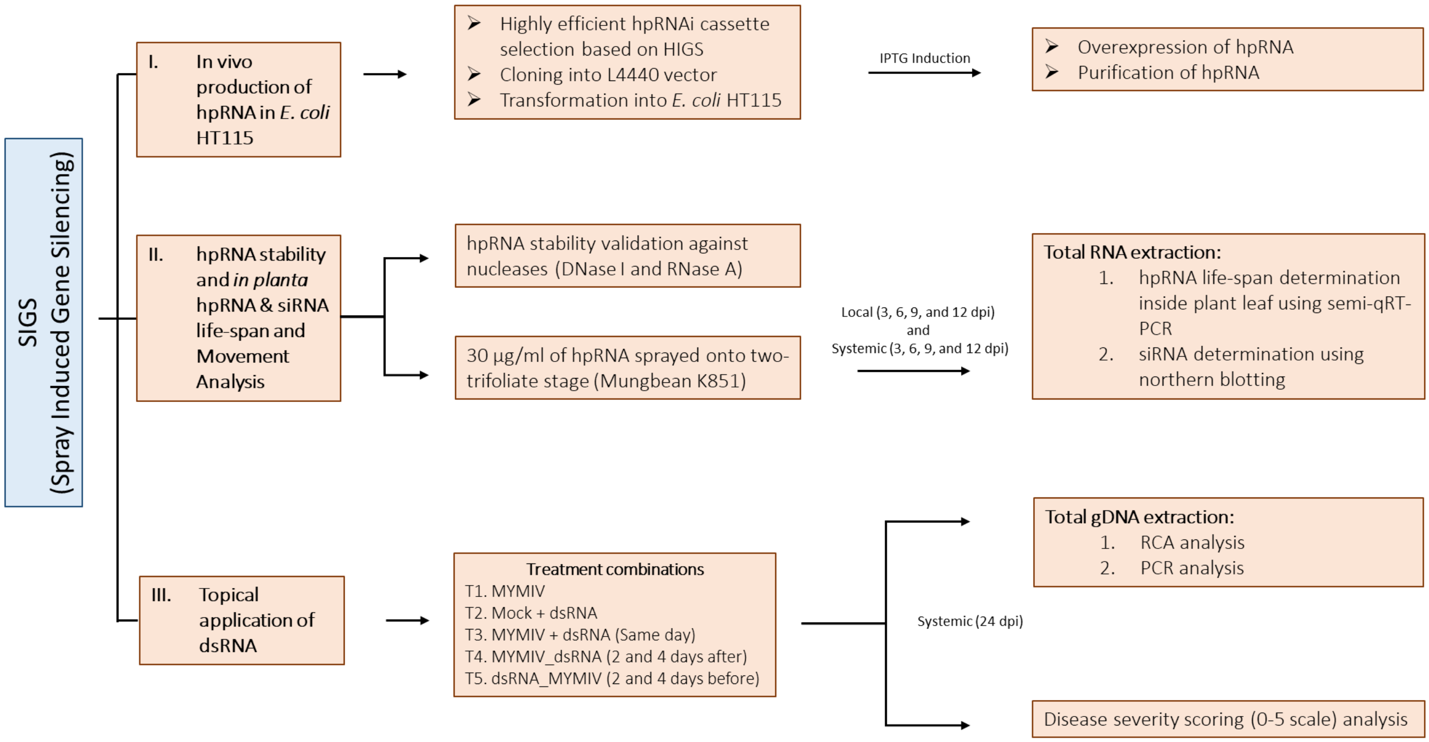


**Fig. S2: Overview of the exogenous application (spray) based resistance methodology in mungbean against Mungbean Yellow Mosaic India Virus (MYMIV).**

The schematic illustrates the key steps and components involved in the experimental approach.

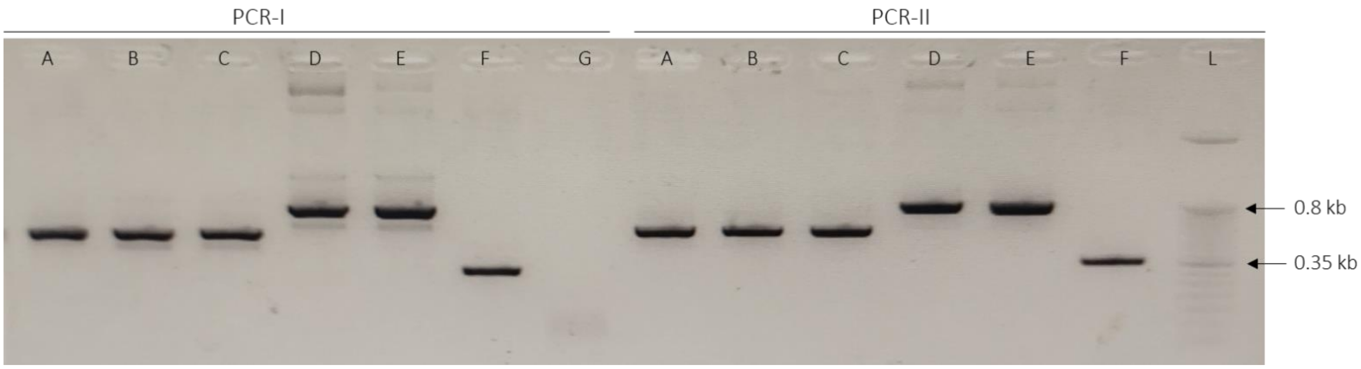


**Fig 1. PCR confirmatory analysis of three hpRNAi constructs in sense and antisense orientation in pKannibal backbone.**

PCR analysis using two different primers set: A: hpTR-1_pKannibal; B: hpTR-2_pKannibal clone I; C: hpTR-2_pKannibal clone II; D: hpTR-1+2_pKannibal clone I; E: hpTR-1+2_pKannibal clone II; F: Empty pKannibal; G: Negative control; L: 50 bp DNA ladder.

**TR-1: AC4/AC1:**

CCTCATCTCCATGTTCTGCTTCAATTCGAAGGAAAGTTGCAAACGAAGAACGAAAGGTTCTTCGACCTGGTTTCCCCAA
CCAGATCAGCACATTACCATCCGAACGTTCAGGCAGCTAAAAGCGCATCAGATGTTAAGTCATACATGGACAAAGACG
GAGACGTCCTTGACCATGGAAGTTTCCAAGTCGATGGCAGATCAGCTAGAGGAGGTAAACAGTCTGCCAACGACGCT
TATGC

**TR-2: AC2/AC3/AC1:**

TTCTCCTCCGTCGATCAAAGCGCAACACAAGGTTGCCAAGAAGCGAGCAATTCGACGCTCTCGAATTGATTTAAGCTGT
GGGTGTAGTTATTACATCCATATCAACTGCCGTAACTATGGATTTTCGCACCGGGGACAACATCACTGCAGCTCAACTC
AGGAATGGCGTCTTTATTTGGGAGGTGCGAAATCCCCTCTCTTTCAAGATCATGCAGCACCGTCAGATTCAAGCAGGG
TCC

**Fig. S3: The nucleotide sequence of TR-1 and TR-2 is depicted.**

Two selected target regions (TRs), TR-1 and TR-2, for RNA interference (RNAi) induction in mungbean against Mungbean Yellow Mosaic India Virus (MYMIV).

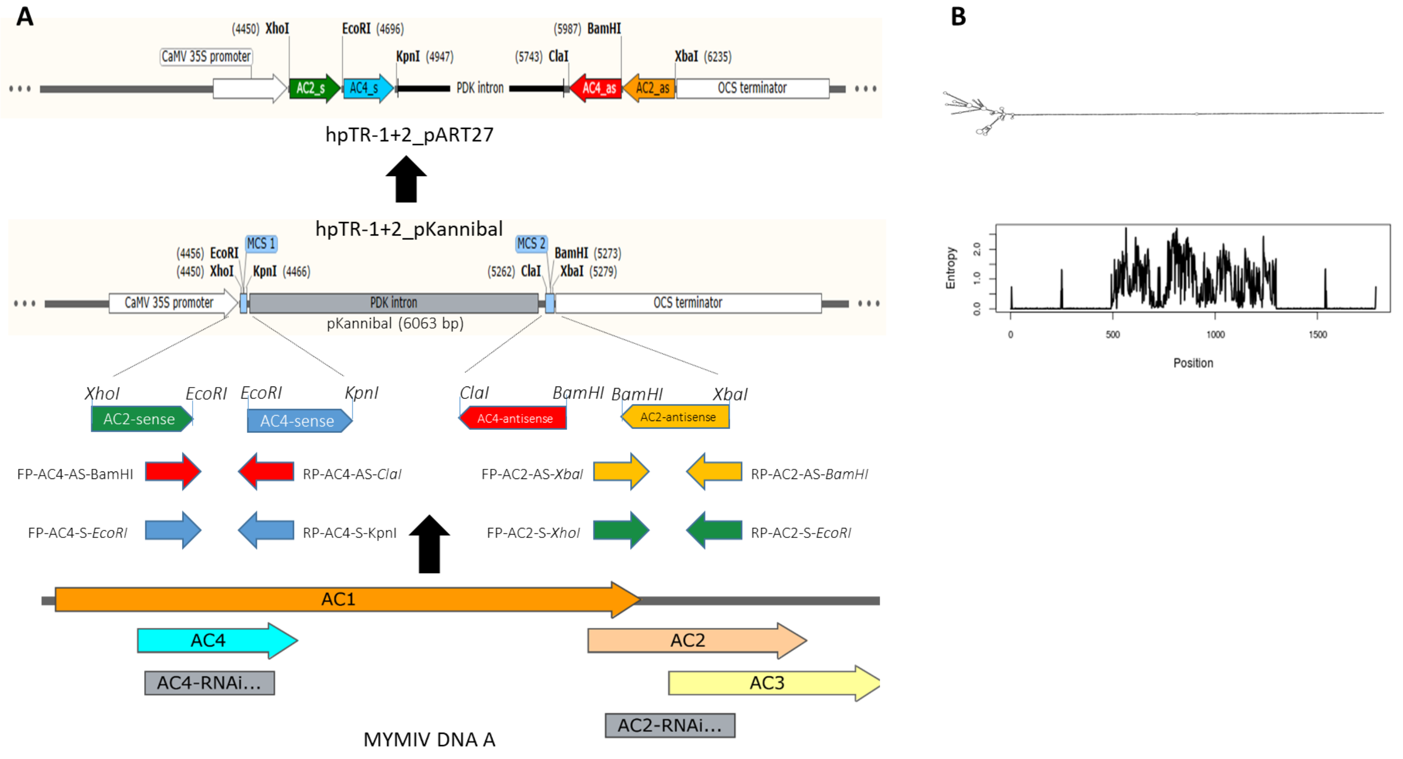


**Fig S4. Schematic representation of hpRNAi clone preparation and hpRNA structure prediction.**

**A)** Construction of hpRNAi cassette, example: hpTR-1+2_pART27 clone preparation is demonstrated; **B**) Schematic representation of predicted dsRNA secondary structure of hpTR-1+2, results have been computed using RNAfold 2.4.18.

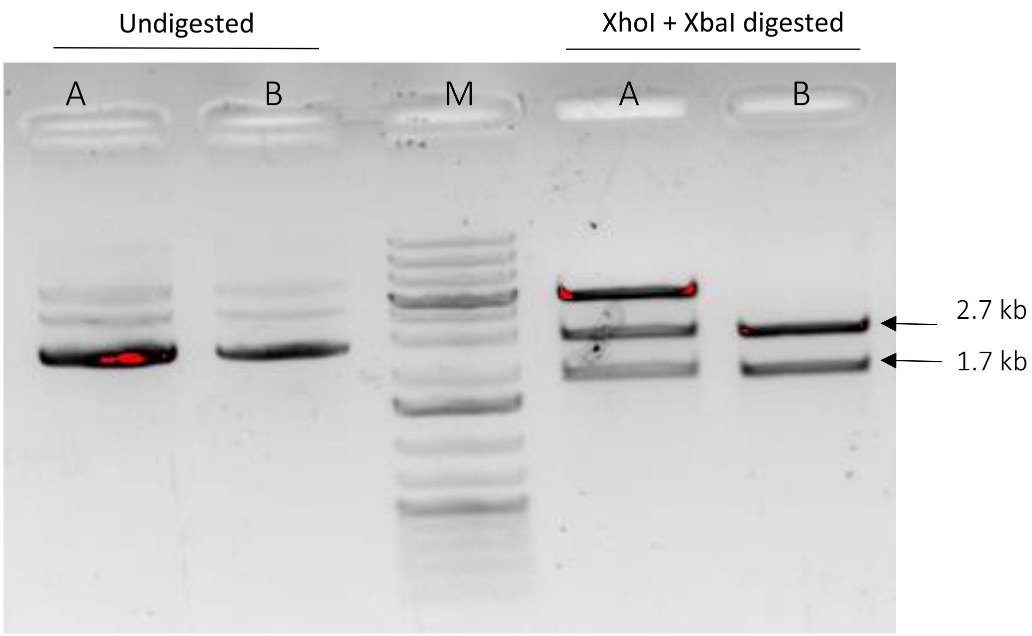


**Fig. S5 Restriction Digestion based confirmation of hpRNA_L4440 recombinant clones:**

Analysis carried out using two enzymes i.e., XhoI and XbaI. A: hpTR-1+2_L4440 clone-I, B: hpTR-1+2_L4440 clone-II, Lane M: 100bp DNA ladder. Appearance of two bands (2.7 kb and 1.7 kb) upon double digestion confirms both sense and antisense fragments are present in L4440 vector backbone.

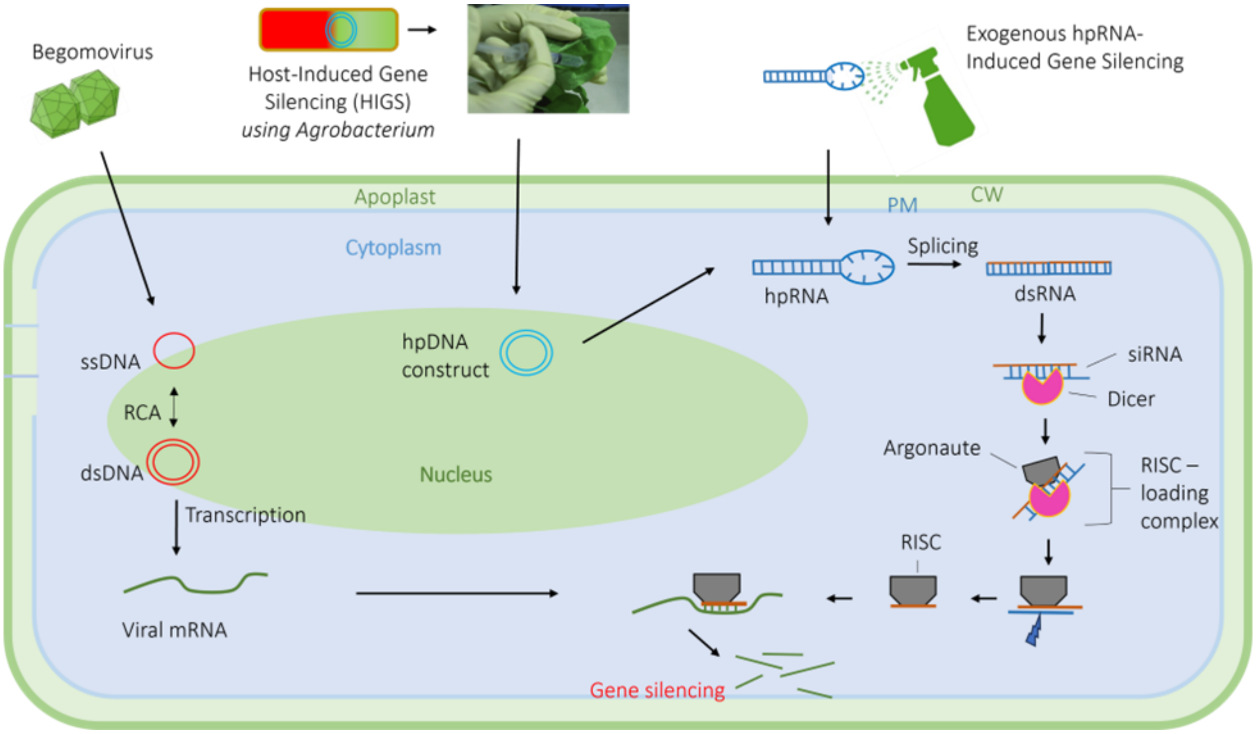


**Fig. S6 The putative mechanism of RNAi activation by HIGS and SIGS.**

The siRNA pathway begins with Dicer's cleavage of double stranded (dsRNA) or hairpin RNA (hpRNA) of exogenous or nuclear origin. The resulting siRNA duplex is loaded onto Argonaute by the RISC-loading complex, which comprises Dicer, a dsRBP protein such as TRBP, and an Argonaute protein. The passenger strand (blue) is cleaved and ejected. The guide strand (brown) remains bound to Argonaute, forming the RISC. The RISC binds to complementary target sequences (green) and silences them via the slicing activity of Argonaute.

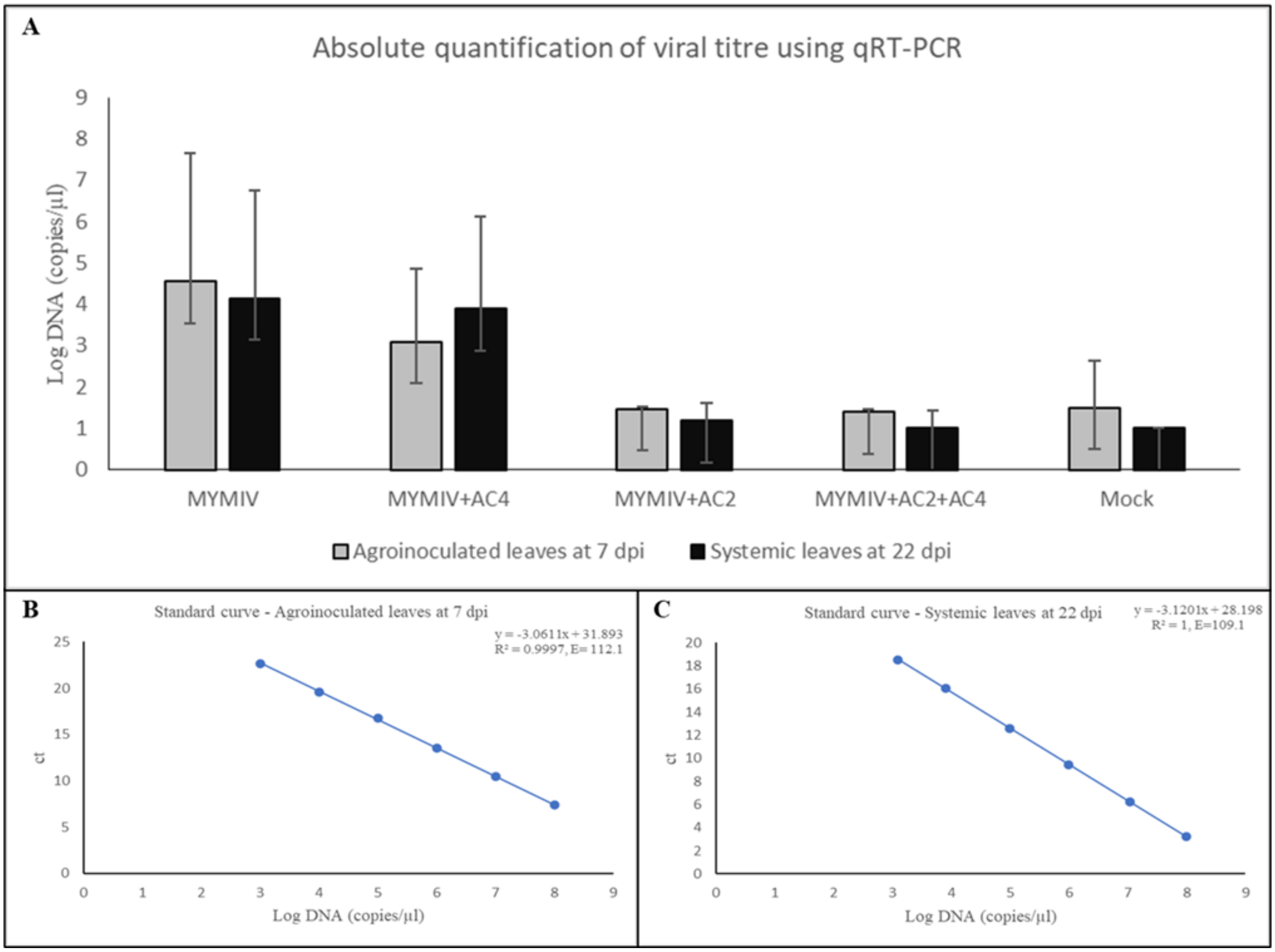


**Fig. S7 Quantification of MYMIV viral titers using quantitative Reverse Transcription Polymerase Chain Reaction (qRT-PCR) in samples from the transient expression assay, which validates the efficacy of three hairpin RNA interference (hpRNAi) constructs.**

The standard curve was generated using a tenfold serial dilution of MYMIV cloned plasmids, with the Ct values plotted against the concentrations of cloned plasmids. B) qRT-PCR standard curve of local leaves 7 dpi, C) qRT-PCR standard curve of systemic leaves 22 dpi.

**Table S1 TR-1 and TR-2 sequence percent Identity. The percent sequence identity analysis of selected RNAi target regions with begomoviruses from various plant using NCBI BLASTn.**

| Virus | Accession | TR-1 Percent Identity | Virus | Accession | TR-2 Percent Identity |
|---|---|---|---|---|---|
| MYMV | OK431081.1 | 100 | MYMV | OK431081.1 | 100 |
| MYMV | JX244176.1 | 99.16 | MYMV | MW736042.1 | 99.16 |
| MYMV | KC911721.1 | 98.74 | MYMV | MW736054.1 | 98.74 |
| MYMV | MN814423.1 | 98.33 | MYMV | AY738104.1 | 98.33 |
| MYMV | OM106038.1 | 97.91 | MYMV | MW792466.1 | 98.32 |
| MYMV | MN698275.1 | 97.49 | MYMV | MN698295.1 | 97.91 |
| MYMV | MN602427.1 | 97.07 | MYMV | KP784665.1 | 99.12 |
| MYMV | MW736048.1 | 96.23 | MYMV | AB017341.1 | 98.68 |
| MYMV | FM242701.1 | 95.82 | MYMV | KP455992.1 | 97.07 |
| MYMV | MW814714.1 | 95.4 | MYMV | MW736046.1 | 98.67 |
| MYMV | OM106037.1 | 94.98 | MYMV | AJ421642.1 | 96.65 |
| MYMV | AM932429.1 | 88.8 | MYMV | OM106037.1 | 95.82 |
| MYMV | KP752088.1 | 88.38 | MYMV | JQ004982.1 | 95.87 |
| MYMV | MW816837.1 | 87.97 | MYMV | AY271896.1 | 95.82 |
| MYMIV | MW814709.1 | 87.92 | MYMV | FM242701.1 | 95.4 |
| MYMIV | MT027035.1 | 87.5 | MYMV | MN885479.1 | 94.98 |
| MYMIV | KP313758.1 | 87.08 | MYMV | D14703.1 | 96.49 |
| MYMIV | OK431079.1 | 87.14 | MYMV | MW792462.1 | 99.02 |
| MYMIV | HF922628.1 | 87.19 | MYMV | MW792465.1 | 99 |
| MYMIV | MT300190.1 | 87.08 | MYMV | MW792463.1 | 98.99 |
| MYMIV | KX363947.1 | 86.25 | MYMIV | AY049772.1 | 91.18 |
| MYMIV | MH255791.1 | 85.83 | MYMIV | MW814710.1 | 90.79 |
| TYLCV | MN842307.1 | 83.9 | MYMIV | MN885468.1 | 98.74 |
| CLCKV | AH013913.2 | 82.08 | MYMIV | MN885463.1 | 98.33 |
| TLCBV | AF428255.1 | 82.08 | HgYMV | MW816839.1 | 93.72 |
| TLCBV | KP164858.1 | 81.67 | HgYMV | KR053204.1 | 92.05 |

| | | | | | |
|---|---|---|---|---|---|
| VBSMV | FN543425.1 | 81.03 | HgYMV | MN698287.1 | 91.63 |
| TLCBV | DQ887537.1 | 80.33 | HgYMV | KP752088.1 | 93.24 |
| TYLCKV | MK946454.1 | 77.69 | HgYMV | AM932425.1 | 91.21 |
| TiCV-2 | MK087038.1 | 78.06 | HgYMV | MW816837.1 | 92.79 |

(Abbrevations: Mungbean yellow mosaic virus (MYMV), Mungbean yellow mosaic India virus (MYMIV), Cotton leaf curl Kokhran virus (CLCKV), Tomato yellow leaf curl virus (TYLCV), Tomato leaf curl Bangalore virus (TLCBV), Velvet bean severe mosaic virus (VBSMV), Tomato yellow leaf curl Kanchanaburi virus (TYLCKV), Tomato interveinal chlorosis virus-2 (TiCV-2), and Horsegram yellow mosaic virus (HgYMV)).

**Table S2 List of primers used to detect MYMIV genome and qRT-PCR based viral titre measurement.**

| | Primer details | Primer code | Sequence (5' to 3') | Length | Amplicon size (bp) |
|---|---|---|---|---|---|
| 1 | *Vigna* radiata, house-keeping gene, i.e. tubulin specific (qRT-PCR) | v.r_tubulin_FP | AACTTATCGATTCCGTCTTGGATG | 24 nt | 184 |
| | | v.r_tubulin_RP | GAAGGGAAAACGGAAAACGTCATCA | 25 nt | |
| 2 | MYMIV DNA-A middle region | Mid_FP | CGTCCATCCATACCTTACCCG | 21 nt | 1210 |
| | | Mid_RP | GTATGCGTCGTTGGCAGATTG | 21 nt | |
| 3 | MYMIV DNA-A , AC1 gene specific (qRT-PCR) | AC1_FP | CTAATAGGTCTATCTGGCCGCG | 22 nt | 137 |
| | | AC1_RP | CGGATATTCACAGAGCCTGTCC | 22 nt | |
